# The m^6^A reader YTHDF2 is a negative regulator for dendrite development and maintenance of retinal ganglion cells

**DOI:** 10.1101/2021.12.07.471547

**Authors:** Fugui Niu, Peng Han, Jian Zhang, Yuanchu She, Lixin Yang, Jun Yu, Mengru Zhuang, Kezhen Tang, Yuwei Shi, Baisheng Yang, Chunqiao Liu, Bo Peng, Sheng-Jian Ji

## Abstract

The precise control of growth and maintenance of the retinal ganglion cell (RGC) dendrite arborization is critical for normal visual functions in mammals. However, the underlying mechanisms remain elusive. Here we find that the m^6^A reader YTHDF2 is highly expressed in the mouse RGCs. Conditional knockout (cKO) of *Ythdf2* in the retina leads to increased RGC dendrite branching, resulting in more synapses in the inner plexiform layer. Interestingly, the *Ythdf2* cKO mice show improved visual acuity compared with control mice. We further demonstrate that *Ythdf2* cKO in the retina protects RGCs from dendrite degeneration caused by the experimental acute glaucoma model. We identify the m^6^A-modified YTHDF2 target transcripts which mediate these effects. This study reveals mechanisms by which YTHDF2 restricts RGC dendrite development and maintenance. YTHDF2 and its target mRNAs might be valuable in developing new treatment approaches for glaucomatous eyes.

**Impact statement:** The m^6^A reader YTHDF2 negatively regulates RGC dendrite branching through destabilizing its m^6^A-modified target mRNAs encoding proteins controlling dendrite development and maintenance. *Ythdf2* cKO improves visual acuity and alleviates acute ocular hypertension-induced glaucoma in mice.

## Introduction

The mammalian retina is an ideal model system to study neuronal development and neural circuit formation. The retinal ganglion cells (RGCs) are the final and only output neurons in the vertebrate retina and their dendrites collect the electrical information concerning the visual signal from all other cells preceding them. One of the major focuses of research in the retina is to understand how RGC dendrite arborization arises during development ***(Prigge and Kay 2018)***. Existing evidences supported that homotypic repulsion controls retinal dendrite patterning ***(Lefebvre et al. 2015)***. However, in mice which had most RGCs genetically eliminated, the dendrite size and shape of remaining RGCs appeared relatively normal ***(Lin et al. 2004)***. Thus, the fact that the dendrites of remaining RGC did not expand to neighboring areas by the remaining RGCs supports the existence of the intrinsic limit for RGC dendrite patterning, which cooperates with the homotypic repulsion to determine the dendrite size of RGCs ***(Lefebvre et al. 2015)***. However, such intrinsic limiting mechanisms remain elusive.

Glaucoma is one of the leading causes for blindness. The major risk factors for glaucoma include increased intraocular tension. Studies have shown that glaucoma causes pathological changes in RGC dendrites before axon degeneration and soma loss were detected in different model animals ***(Weber et al. 1998; Shou et al. 2003; Morgan et al. 2006)***. Thus, elucidation of mechanisms governing RGC dendrite arbor maintenance bears clinical significance.

*N*^6^-methyladenosine (m^6^A) is the most widely distributed and extensively studied internal modification in mRNA ***(Dominissini et al. 2012; Meyer et al. 2012; Nachtergaele and He 2018)***. m^6^A modification has been shown to regulate brain development and functions in the nervous system ***(Livneh et al. 2020; Yu et al. 2021)***. By effectors, most of these studies have focused on its demethylases (“m^6^A erasers”) and methyltransferases (“m^6^A writers”). Since the fate of m^6^A-modified transcripts are decoded by the m^6^A binding proteins (“m^6^A readers”), how the readers mediate these functions and what are their neural target mRNAs remain to be elucidated. In addition, more precisely controlled spatial-temporal ablation of the m^6^A readers instead of null knockout is required to elucidate their functions and mechanisms in nervous system.

In this study, we identified an m^6^A-dependent intrinsic limiting mechanism for RGC dendrite arborization and maintenance. Conditional knockout of the m^6^A reader YTHDF2 in the developing mouse retina increases RGC dendrite branching and improves visual acuity. YTHDF2 also mediates acute ocular hypertension (AOH)-induced RGC degeneration, the experiment model for glaucoma, and *Ythdf2* cKO in the retina alleviates AOH-induced RGC dendrite shrinking and neuronal loss. The regulation of RGC dendrite development and maintenance by YTHDF2 is mediated by two distinct groups of m^6^A-modified target mRNAs which encode proteins that promote dendrite arborization during development and maintain dendrite tree during injury, respectively. Therefore, our study reveals mechanisms by which YTHDF2 restricts RGC dendrite development and maintenance, which sheds light on developing new treatment approaches for glaucomatous eyes.

## Results

### Knockdown of YTHDF2 leads to a robust increase of RGC dendrite branching

To examine whether m^6^A modification and its reader proteins play a role in the dendrite development, we utilized the retina as the model system. We first checked their expression patterns in the developing mouse retina. Immunostaining with a widely used m^6^A antibody demonstrated that RGCs had high m^6^A modification levels (*Figure 1—figure supplement 1A*). Consistent with the m^6^A distribution, the m^6^A reader YTHDF2 is highly expressed in RGCs (*Figure 1A*; *Figure 1—figure supplement 1A*). Conversely, the expression of YTHDF2 in other layers of the retina is much lower (*Figure 1A*; *Figure 1—figure supplement 1B*). Another two m^6^A readers YTHDF1 and YTHDF3 show similar expression patterns (*Figure 1—figure supplement 1C,D*). The strong expression of YTHDFs and high level of m^6^A modification in RGCs suggest that the m^6^A reader YTHDFs might play roles in RGC development. We dissected and dissociated the retinal cells and cultured in vitro. We generated lenti viral shRNAs against YTHDFs, which showed similarly efficient knockdown (KD) of YTHDFs in RGC cultures in vitro (*Figure 1B*; *Figure 1—figure supplement 1E,F*). In these YTHDF-deficient RGC cultures, the first and most obvious phenotype that we observed is the robust increase of dendrite branching of cultured RGCs treated by *shYthdf2* (*Figure 1C,D*; *Figure 1—figure supplement 1G,H*). In contrast, the dendrite branching of RGCs with YTHDF1 KD using *shYthdf1* was not significantly different from control shRNA (*Figure 1—figure supplement 1I*), while YTHDF3 KD using *shYthdf3* caused a slight (statistically significant in several Sholl radii) decrease of RGC dendrite branching compared with control shRNA (*Figure 1—figure supplement 1J*). These results suggest that the m^6^A reader YTHDF2 might play an important role in controlling dendrite branching of RGCs.

**Figure 1.**
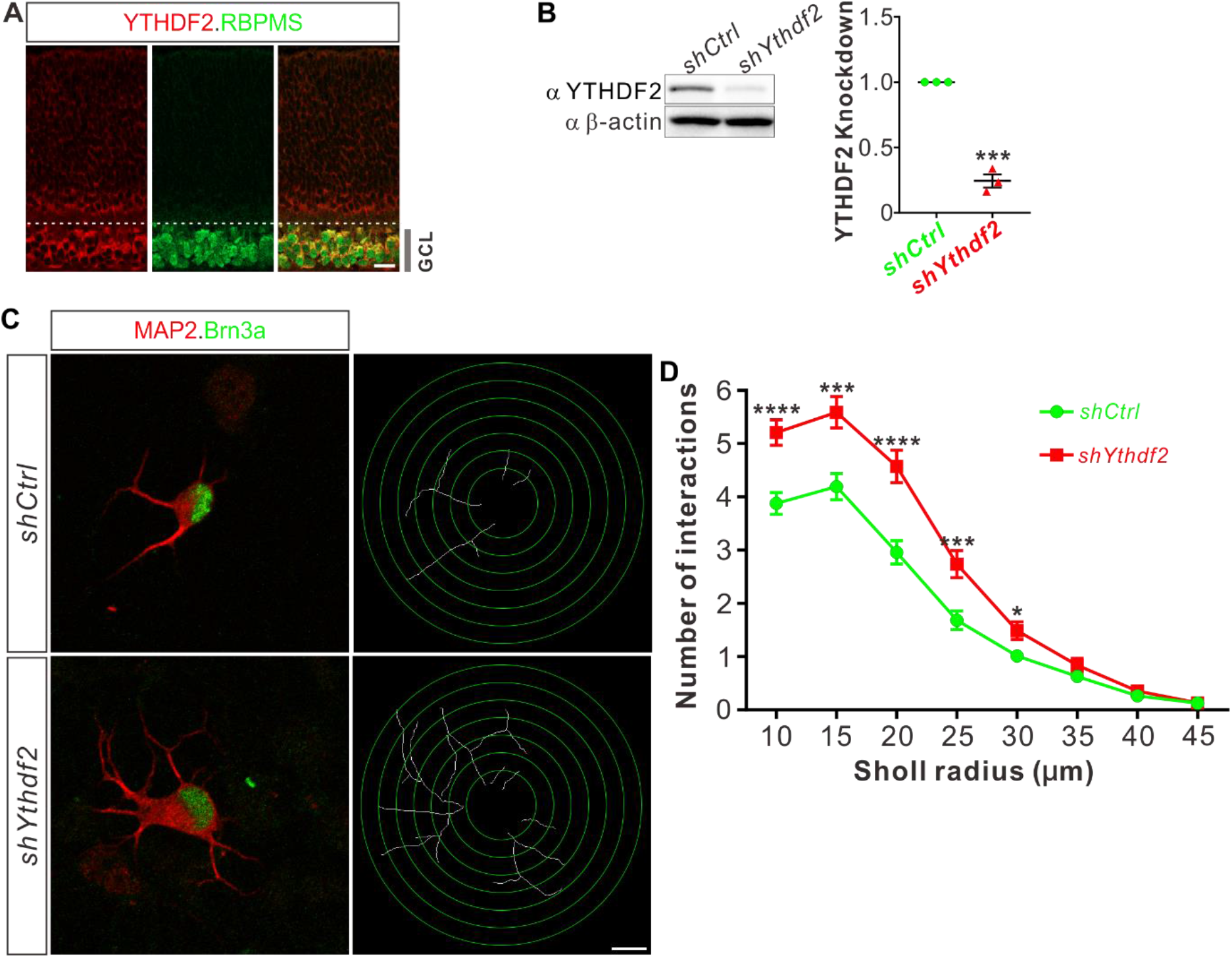
Knockdown of YTHDF2 leads to a robust increase of RGC dendrite branching. (**A**) Representative confocal images showing high expression of YTHDF2 in RGCs (marked by RBPMS) in P0 retina. Note that all RGCS marked by the pan-RGC marker RBPMS express YTHDF2 while all YTHDF2-expressing cells are RBPMS^+^ RGCs. GCL, ganglion cell layer. Scale bars: 20 μm. (**B**) Western blotting (WB) confirming efficient knockdown (KD) of YTHDF2 in cultured RGCs using *shYthdf2*. Data of WB quantification are mean ± SEM and are represented as dot plots: \*\*\**p* = 0.00012 (*n* = 3 replicates); by unpaired Student’s *t* test. (**C**) Examination of RGC dendrite development after YTHDF2 KD. As shown, significantly increased branching of dendrites marked by MAP2 immunofluorescence was observed in cultured RGCs marked by Brn3a. Dendrite traces were drawn for the corresponding RGCs. Scale bar: 10 μm. (**D**) Quantification of dendrite branching (**C**) using Sholl analysis. As shown, numbers of interactions are significantly greater in *shYthdf2* groups (*n* = 68 RGCs) than *shCtrl* groups (*n* = 72 RGCs) in Sholl radii between 10-30 μm. Data are mean ± SEM. \*\*\*\**p* = 4.32E-05 (10 μm), \*\*\**p* = 0.00038 (15 μm), \*\*\*\**p* = 2.85E-05 (20 μm), \*\*\**p* = 0.00084 (25 μm), \**p* = 0.020 (30 μm), by unpaired Student’s *t* test.

### Conditional knockout of *Ythdf2* in the retina increases RGC dendrite branching in vivo without disturbing sublaminar targeting

To further explore whether YTHDF2 physiologically regulates RGC dendrite branching in vivo, we generated *Ythdf2* conditional knockout (*Ythdf2* cKO) mouse (*Figure 2A*). We used the *Six3-cre* mouse line ***(Furuta et al. 2000)***, which has been widely used in the field to generate retina-specific knockouts ***(Lefebvre et al. 2012; Riccomagno et al. 2014; Sapkota et al. 2014; Krishnaswamy et al. 2015)***. YTHDF2 expression is efficiently eliminated in the *Ythdf2* cKO retina compared with their littermate controls at E12.5 (*Figure 2—figure supplement 1A*) and E15.5 (*Figure 2B*). Retina progenitors, amacrine cells, bipolar cells, photoreceptors, or horizontal cells were not affected in *Ythdf2* cKO retina (*Figure 2—figure supplement 1B-L*), suggesting that YTHDF2 is not involved in the generation or development of these cells. This is in line with the low YTHDF2 expression in these cells. The RGC number or density was not affected in the *Ythdf2* cKO retina (*Figure 2C,D*), demonstrating that *Ythdf2* knockout does not disturb RGC neurogenesis. We then cultured RGCs from the *Ythdf2* cKO retina. The dendrite branching of *Ythdf2* cKO RGCs was significantly increased compared with littermate controls (*Figure 2E,F*). RGCs include over 40 subtypes ***(Sanes and Masland 2015; Baden et al. 2016)***. We thus examined the RGC dendrite branching within different subtypes. One of the RGC subgroups responds preferentially to movement in particular directions and is named the ON-OFF directionally selective RGCs (ooDSGCs). Expression of CART (cocaine- and amphetamine-regulated transcript), a neuropeptide, distinguishes ooDSGCs from other RGCs ***(Kay et al. 2011)***. The dendrite branching of ooDSGCs marked by CART/Brn3a co-staining in *Ythdf2* cKO retinal cultures also increased compared with control (*Figure 2G,H*). These data further confirm that the m^6^A reader YTHDF2 regulates dendrite branching of RGCs.

**Figure 2.**
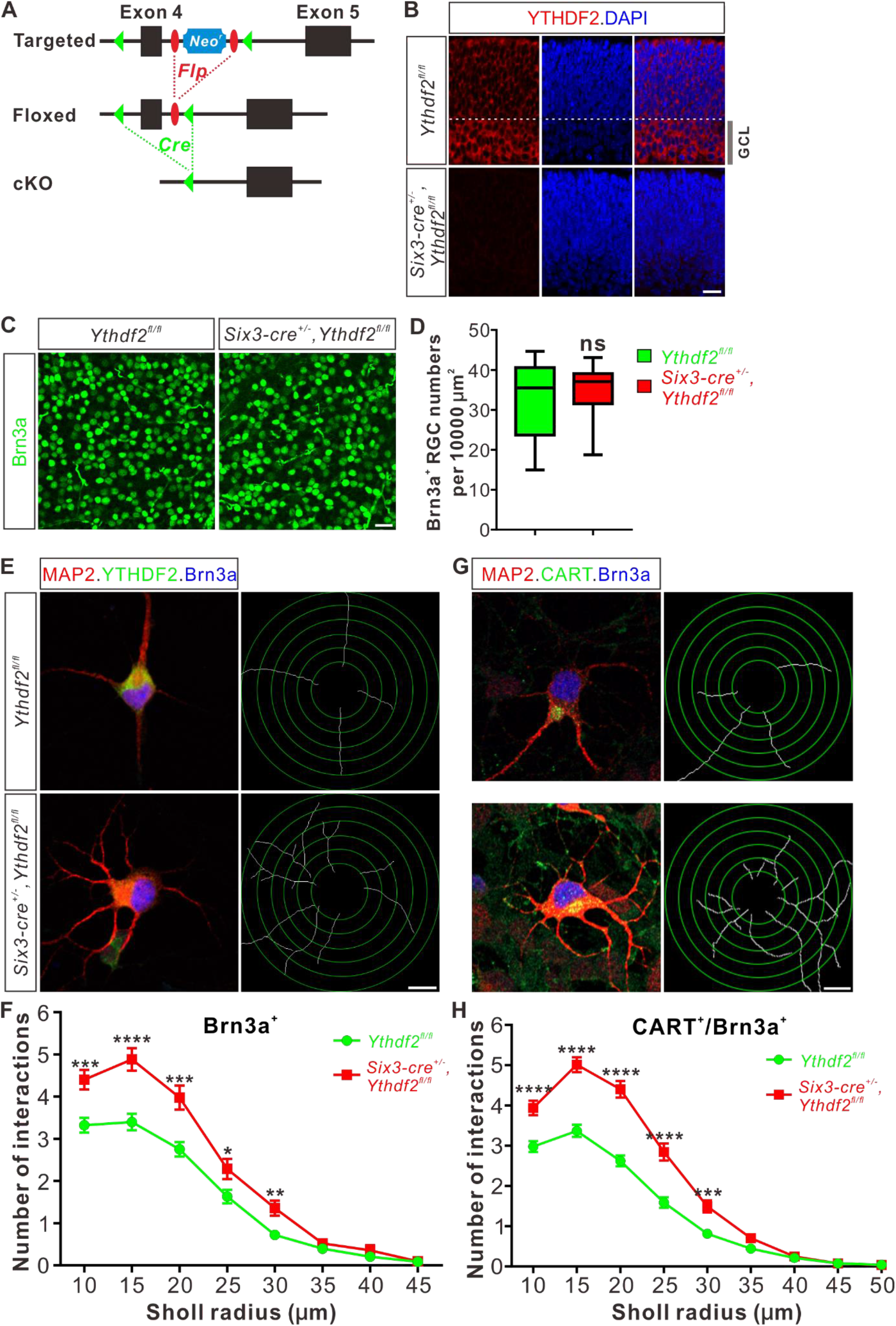
Dendrite branching is dramatically increased in cultured RGCs from *Ythdf2* cKO. (A) Schematic drawings of the genetic deletion strategy for *Ythdf2*. Exon 4 which contains YTH domain-coding sequence is deleted after Cre-mediated recombination. (**B**) Depletion of YTHDF2 protein in retina of *Six3-cre^+/-^;Ythdf2^fl/fl^* cKO mice. Anti-YTHDF2 immunostaining of E15.5 retina vertical sections confirmed cKO of YTHDF2 protein, compared with *Ythdf2^fl/fl^* littermate controls. Scale bar: 20 μm. (**C, D**) RGC neurogenesis not affected in the *Ythdf2* cKO retina. Wholemount immunostaining using a Brn3a antibody was carried out in P20 retina (**C**). Numbers of Brn3a^+^ RGC per 10000 μm^2^ of retina were quantified and showed no difference between the *Ythdf2* cKO and their littermate controls (**D**). *n* = 12 confocal fields for each genotype. Data are represented as box and whisker plots: ns, not significant (*p* = 0.79); by unpaired Student’s *t* test. Scale bar: 25 μm. (**E**) Examination of RGC dendrite development in *Ythdf2* cKO RGCs. As shown, knockout of YTHDF2 was confirmed by YTHDF2 IF (green). Significantly increased branching of dendrites marked by MAP2 IF (red) was observed in cultured RGCs from the *Ythdf2* cKO retina compared with their littermate controls. Dendrite traces were drawn for the corresponding RGCs. Scale bar: 10 μm. (**F**) Quantification of RGC dendrite branching (**E**) using Sholl analysis. Data are mean ± SEM. Numbers of interactions are significantly greater in *Six3-cre^+/-^,Ythdf2^fl/fl^* groups (*n* = 68 RGCs) than *Ythdf2^fl/fl^* groups (*n* = 42 RGCs) in Sholl radii between 10-30 μm: \*\*\**p* = 0.00030 (10 μm), \*\*\*\**p* = 1.19E-05 (15 μm), \*\*\**p* = 0.00018 (20 μm), \**p* = 0.021 (25 μm), \*\**p* = 0.0022 (30 μm), by unpaired Student’s *t* test. (**G**) Examination of CART^+^ RGC dendrite development in *Ythdf2* cKO RGCs. Cultured CART^+^ RGCs from the *Ythdf2* cKO retina have significantly increased branching of dendrites marked by MAP2 IF (red) compared with their littermate controls. Dendrite traces were drawn for the corresponding RGCs. Scale bar: 10 μm. (**H**) Quantification of CART^+^ RGC dendrite branching (**G**) using Sholl analysis. Data are mean ± SEM. Numbers of interactions are significantly greater in *Six3-cre^+/-^,Ythdf2^fl/fl^* groups (*n* = 77 RGCs) than *Ythdf2^fl/fl^* groups (*n* = 90 RGCs) in Sholl radii between 10-30 μm: \*\*\*\**p* = 3.17E-05 (10 μm), \*\*\*\**p* = 6.50E-11 (15 μm), \*\*\*\**p* = 5.14E-12 (20 μm), \*\*\*\**p* = 5.00E-07 (25 μm), \*\*\**p* = 0.00020 (30 μm), by unpaired Student’s *t* test.

Next, we wanted to confirm this phenotype in vivo by checking specific RGC subtypes. Intravitreal injection of an AAV reporter expressing ZsGreen visualized the dendrite morphology of ooDSGCs marked by CART immunostaining (*Figure 3A*). ooDSGCs showed dramatically increased dendrite branching in *Ythdf2* cKO retina compared with control retina by Sholl analysis (*Figure 3A,B*). The intrinsically photosensitive RGCs (ipRGCs) are unique and melanopsin-expressing cells, which exhibit an intrinsic sensitivity to light ***(Hattar et al. 2002)***. We analyzed the morphology of ipRGCs visualized by wholemount immunostaining of melanopsin and found that the dendrite branching of ipRGCs was significantly increased in the *Ythdf2* cKO retina (*Figure 3C,D*; *Figure 3—figure supplement 1A-E*). A similar trend was observed in the SMI-32^+^ αRGCs (*Figure 3E,F*). These results strongly indicate that the m^6^A reader YTHDF2 negatively regulates RGC dendrite branching in vivo and *Ythdf2* cKO promotes RGC dendrite arborization.

**Figure 3.**
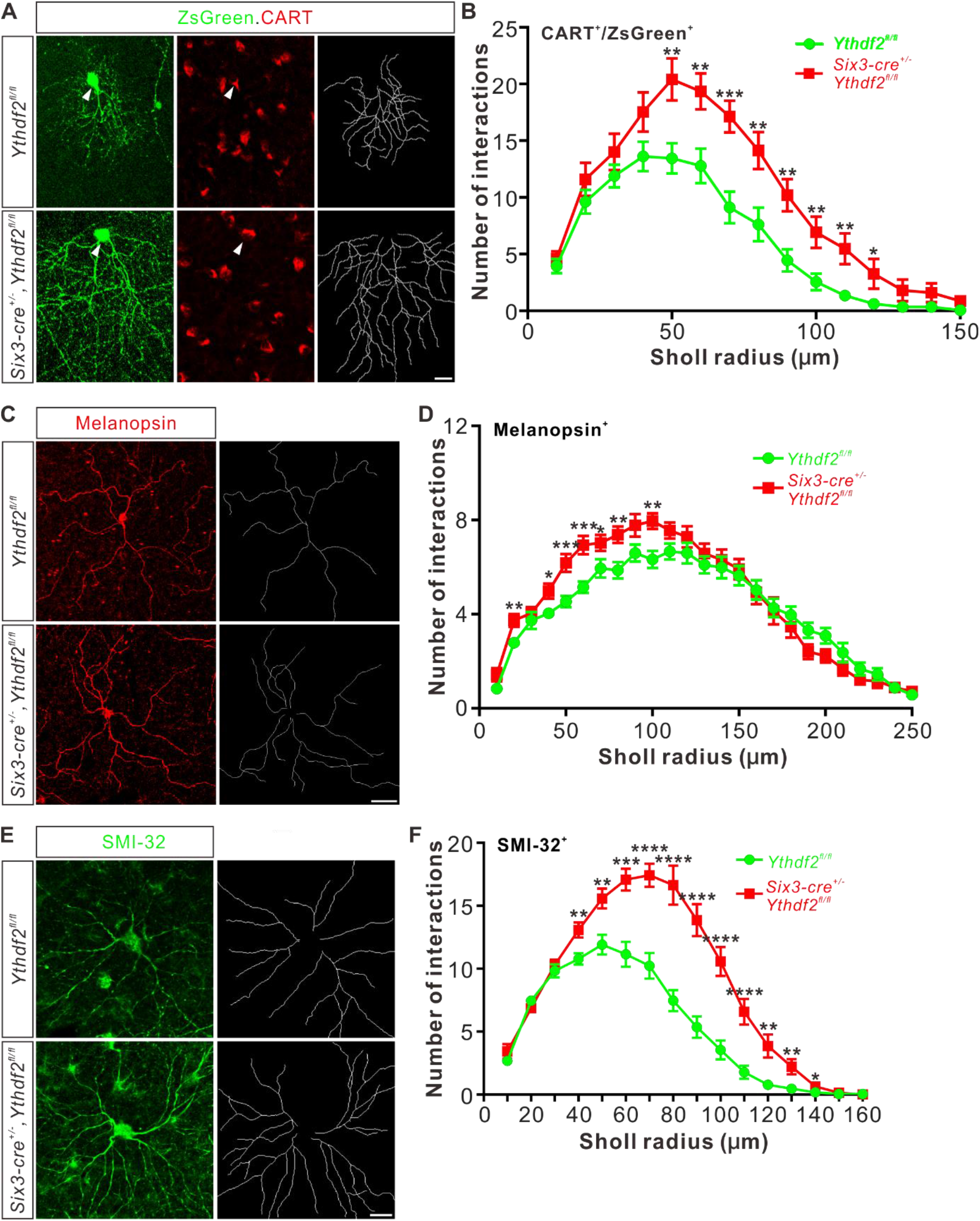
Dendrite branching of specific RGC subtypes increases in *Ythdf2* cKO in vivo. (**A**) Co-labeling of ooDSGCs by AAV-ZsGreen and CART IF in vivo. Intravitreal injection of AAV expressing ZsGreen reporter was performed at P17 and retinas were collected at P27. The white arrowheads indicate ooDSGCs co-labeled by ZsGreen and CART IF, which show dramatically increased dendrite branching in *Ythdf2* cKO compared with control. Dendrite traces were drawn for the corresponding RGCs shown. Scale bar: 20 μm. (**B**) Quantification of dendrite branching of ZsGreen^+^/CART^+^ ooDSGCs (**A**) using Sholl analysis. Data are mean ± SEM. Numbers of interactions are significantly greater in *Six3-cre^+/-^,Ythdf2^fl/fl^* groups (*n* = 15 RGCs) than *Ythdf2^fl/fl^* groups (*n* = 18 RGCs) in Sholl radii between 50-120 μm: \*\**p* = 0.0041 (50 μm), \*\**p* = 0.0059 (60 μm), \*\*\**p* = 0.00036 (70 μm), \*\**p* = 0.0058 (80 μm), \*\**p* = 0.0018 (90 μm), \*\**p* = 0.0064 (100 μm), \*\**p* = 0.0045 (110 μm), \**p* = 0.040 (120 μm), by unpaired Student’s *t* test. (**C**) Dendrites of ipRGCs visualized by wholemount immunostaining of P20 retina using a melanopsin antibody in vivo. Dendrite traces were drawn for the corresponding RGCs shown. Scale bar: 50 μm. (**D**) Quantification of dendrite branching of melanopsin^+^ ipRGCs (**C**) using Sholl analysis. Data are mean ± SEM. Numbers of interactions are significantly greater in *Six3-cre^+/-^,Ythdf2^fl/fl^* groups (*n* = 18 RGCs) than *Ythdf2^fl/fl^* groups (*n* = 21 RGCs) in Sholl radii between 20-100 μm: \*\**p* = 0.0083 (20 μm), \**p* = 0.018 (40 μm), \*\*\**p* = 0.00068 (50 μm), \*\*\**p* = 0.00027 (60 μm), \**p* = 0.048 (70 μm), \*\**p* = 0.0048 (80 μm), \*\**p* = 0.0023 (100 μm), by unpaired Student’s *t* test. (**E**) Dendrites of αRGCs visualized by wholemount immunostaining of P20 retina using a SMI-32 antibody in vivo. Dendrite traces were drawn for the corresponding RGCs shown. Scale bar: 20 μm. (**F**) Quantification of dendrite branching of SMI-32^+^ αRGCs (**E**) using Sholl analysis. Data are mean ± SEM. Numbers of interactions are significantly greater in *Six3-cre^+/-^,Ythdf2^fl/fl^* groups (*n* = 14 RGCs) than *Ythdf2^fl/fl^* groups (*n* = 22 RGCs) in Sholl radii between 40-140 μm: \*\**p* = 0.0044 (40 μm), \*\**p* = 0.0035 (50 μm), \*\*\**p* = 0.00021 (60 μm), \*\*\*\**p* = 2.63E-05 (70 μm), \*\*\*\**p* = 2.38E-06 (80 μm), \*\*\*\**p* = 1.68E-06 (90 μm), \*\*\*\**p* = 6.76E-06 (100 μm), \*\*\*\**p* = 5.72E-05 (110 μm), \*\**p* = 0.0011 (120 μm), \*\**p* = 0.0032 (130 μm), \**p* = 0.047 (140 μm), by unpaired Student’s *t* test.

In the retina, RGCs target their dendrites in different sublaminae of the inner plexiform layer (IPL). Since the IPL sublaminar targeting of RGC dendrites is critical for normal visual functions, we wondered whether the increased dendrite branching caused by *Ythdf2* cKO was also accompanied by altered sublaminar patterning of RGC dendrites. We used a *Thy1-GFP* reporter (line O) which labels a few RGCs ***(Feng et al. 2000)***. As shown in *Figure 3—figure supplement 1F,G*, GFP intensity is generally higher in IPL of the *Ythdf2* cKO retina compared with their littermate controls, which further proves the increased RGC dendrite branching and density. However, the sublaminar pattern of GFP signals looks similar between cKO and littermate control (*Figure 3—figure supplement 1F,G*). Sublaminar dendrite patterning of the ipRGC subtype visualized by immunostaining of melanopsin also demonstrated the similar phenotype (*Figure 3—figure supplement 1H,I*). These data suggest that YTHDF2 has a general control of RGC dendrite branching but has no striking effect on the sublaminar targeting of RGC dendrite. These results are consistent with the previous findings that the RGC dendrite targeting is determined genetically and several transcription factors controlling laminar choice have been identified in RGCs and amacrine cells ***(Cherry et al. 2011; Kay et al. 2011; Lefebvre et al. 2015; Liu et al. 2018)***.

### IPL of *Ythdf2* cKO retina is thicker and has more synapses

The increased dendrite branching of RGCs further prompted us to check whether *Ythdf2* cKO changes IPL development. Immunostaining of P6 retina vertical sections using a MAP2 antibody demonstrated that IPL thickness significantly increased in *Ythdf2* cKO retina (*Figure 4A,B*). As a control, the thicknesses of other retinal layers showed no difference between the *Ythdf2* cKO and control mice (*Figure 4—figure supplement 1A-D*). Quantification of MAP2 IF intensity in IPL suggested that the IPL of *Ythdf2* cKO retina became denser with dendrites (*Figure 4A,C*). These results suggest that the increased dendrite branching results in a thicker and denser IPL in the *Ythdf2* cKO retina.

**Figure 4.**
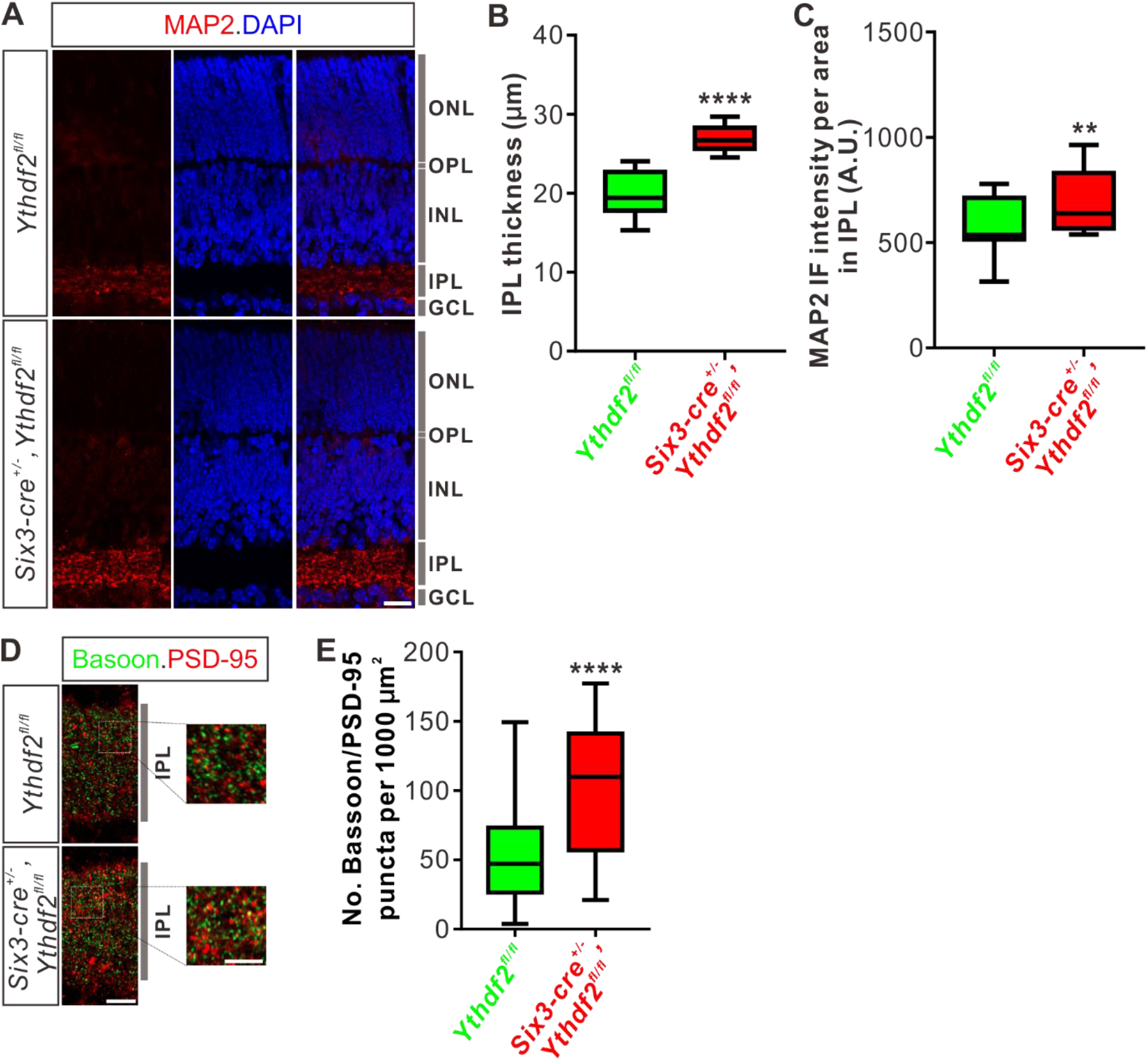
IPL of the *Ythdf2* cKO retina is thicker and has more synapses. (**A**) Cross-sections of P6 *Six3-cre^+/-^,Ythdf2^fl/fl^* retina showing increased IPL thickness by MAP2 staining compared with littermate control. ONL, outer nuclear layer; OPL, outer plexiform layer; INL, inner nuclear layer; IPL, inner plexiform layer; GCL, granule cell layer. Scale bar: 20 μm. (**B, C**) Quantification showing increased IPL thickness and MAP2 IF intensity per area in IPL of the *Ythdf2* cKO retina (**A**). Quantification data are represented as box and whisker plots: \*\*\**p* = 1.28E-07 for **B** (*n* = 12 sections for each genotype), by unpaired Student’s *t* test; \*\**p* = 0.0045 for **C** (*n* = 12 sections for each genotype), by paired Student’s *t* test. (**D, E**) Representative confocal images showing the excitatory synapses labeled by co-localization of Bassoon (presynaptic) and PSD-95 (postsynaptic) in the IPL of P30 retina (**D**). There are significantly more synapses in the *Ythdf2* cKO IPL compared with control. Quantification data are represented as box and whisker plots (**E**): *n* = 47 confocal fields for *Ythdf2^fl/fl^*, *n* = 23 confocal fields for *Six3-cre^+/-^, Ythdf2^fl/fl^*; \*\**p* = 1.63E-05; by unpaired Student’s *t* test. Scale bars: 10 μm (**D**) and 5 μm (inset in **D**).

The inner plexiform layer (IPL) of retina is concentrated with synaptic connections, which contain synapses among and between bipolar-amacrine-ganglion cells. The increased RGC dendrite branching and denser IPL in the *Ythdf2* cKO retina prompted us to wonder whether there are changes in synaptic connections in IPL. We used co-staining of the presynaptic marker Bassoon and the postsynaptic marker PSD-95 to count the colocalization puncta of Bassoon^+^/PSD-95^+^. We found that the numbers of Bassoon^+^/PSD-95^+^ excitatory synapses in IPL of *Ythdf2* cKO retina are significantly larger than that of control retina (*Figure 4D,E*). As a control, the numbers of the excitatory ribbon synapses marked by the colocalization of Bassoon^+^/PSD-95^+^ in OPL (outer plexiform layer) show no difference between *Ythdf2* cKO and control retinas (*Figure 4—figure supplement 1E,F*).

All these data verify that the IPL of *Ythdf2* cKO retina is thicker and has more synapses.

### Visual acuity is improved for the *Ythdf2* cKO mice

The features of RGC dendrites, including their size, shape, arborization pattern and localization, influence the amount and type of synaptic inputs that RGCs receive, which in turn determine how RGCs respond to specific visual stimuli such as the direction of motion ***(Liu and Sanes 2017)***. The increased dendrite branching, the thicker and denser IPL, and the more synapses in the IPL inspired us to further explore whether the visual responses of the *Ythdf2* cKO mice were changed or not. *Ythdf2* cKO mice looked normal and had similar body weight and size compared with control mice for either sex (male in *Figure 5A,B*; female in *Figure 5C,D*). The generally normal development of *Ythdf2* cKO mice is consistent with the specific and limited expression of *Six3-cre* in retina (*Figure 5—figure supplement 1A*), and only sparse spots in ventral forebrain (*Figure 5—figure supplement 1B*) ***(Furuta et al. 2000)***. We used an optomotor response (OMR)-based assay ***(Prusky et al. 2004; Umino et al. 2008; Shi et al. 2018)*** to monitor visual functions of *Ythdf2* cKO mice (*Figure 5E*). Surprisingly, the *Ythdf2* cKO mice showed modestly improved visual acuity compared with the control mice, measuring spatial frequency threshold as 0.45 ± 0.0043 c/deg (cycle per degree) and 0.43 ± 0.0085 c/deg, respectively (*Figure 5F*, male mice). Similar phenotype was observed in female mice (*Figure 5G*). These results suggest that the visual acuity is modestly improved in the *Ythdf2* cKO mice.

**Figure 5.**
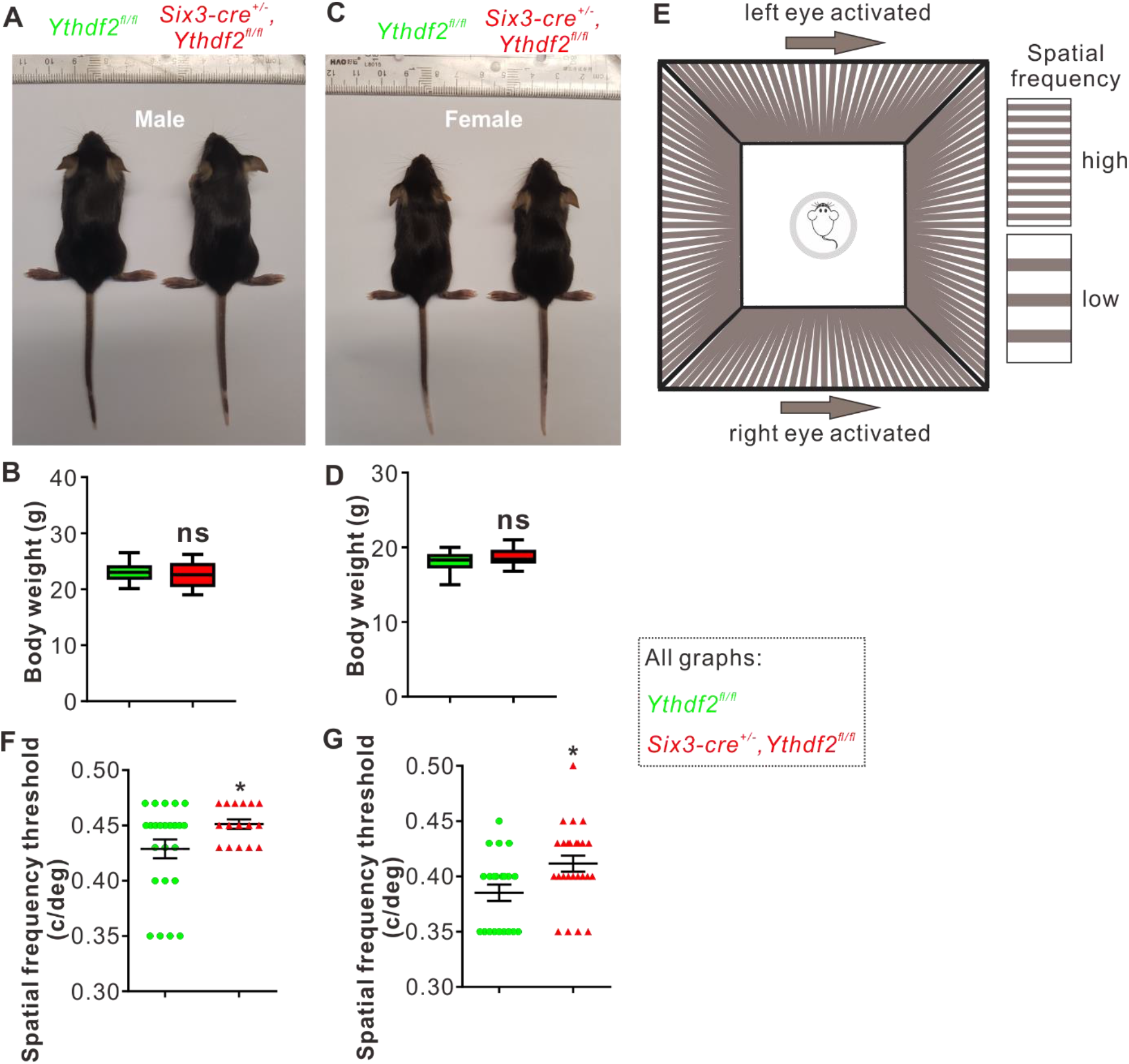
Visual acuity is improved for the *Ythdf2* cKO mice. (**A-D**) Six3-Cre-mediated *Ythdf2* cKO showing normal animal development and body weight (male in **A**, female in **C**). Quantification data of body weight (**B, D**) are represented as box and whisker plots: *p* = 0.41 in **B** (male, *n* = 24 for control, *n* = 18 for cKO); *p* = 0.08 in **D** (female, *n* = 23 for control, *n* = 25 for cKO); ns, not significant; by unpaired Student’s *t* test. (**E**) The setup of optomotor response assay is illustrated by schematic drawing. (**F, G**) Optomotor response assay demonstrating improved visual acuity in the *Ythdf2* cKO mice. Quantification data are mean ± SEM: \**p* = 0.048 in **F** (male, *n* = 24 control, *n* = 16 cKO); \**p* = 0.015 in **G** (female, *n* = 21 control, *n* = 25 cKO); by unpaired Student’s *t* test.

This phenotype is most likely attributed to the increased RGC dendrite branching and thicker and denser IPL with more synapses because all other parts and processes of retina are not affected except RGC dendrite in the *Ythdf2* cKO mediated by *Six3-cre* (*Figure 2—figure supplement 1 and Figure 4—figure supplement 1*). The eyes and optic fibers also showed no difference between *Ythdf2* cKO and control mice (*Figure 5—figure supplement 1C-E*). We further checked the targeting of optic nerves to the brain by anterograde labeling with cholera toxin subunit B (CTB) and found no difference of retinogeniculate or retinocollicular projections between *Ythdf2* cKO and control mice (*Figure 5—figure supplement 1F,G*), suggesting the guidance and central targeting of RGC axons are not affected in the *Ythdf2* cKO.

### YTHDF2 target mRNA were identified with transcriptomic and proteomic analysis

Next, we continued to explore the underlying molecular mechanisms of the effects on dendrite branching caused by *Ythdf2* cKO in the retina. First, we wanted to know what transcripts YTHDF2 recognizes and binds. We carried out anti-YTHDF2 RNA immunoprecipitation (RIP) in the retina followed by RNA sequencing of the elute (RIP-Seq). Two biological replicates of anti-YTHDF2 RIP-Seq identified 1638 transcripts (*Supplementary file 1*). Functional annotation of YTHDF2 RIP targets revealed significant enrichment in Cellular Component terms such as neuron part and neuron projection, and Biological Process terms such as cellular component organization and neuron projection development. We further zoomed in to check neural terms in Cellular Component (*Figure 6A*) and Biological Process (*Figure 6B*). We found that substantial numbers of YTHDF2 target transcripts are involved in cytoskeleton, dendrite and their organization and development (*Figure 6A,B*), which is consistent with the dendrite branching phenotype observed in the *Ythdf2* cKO retina.

**Figure 6.**
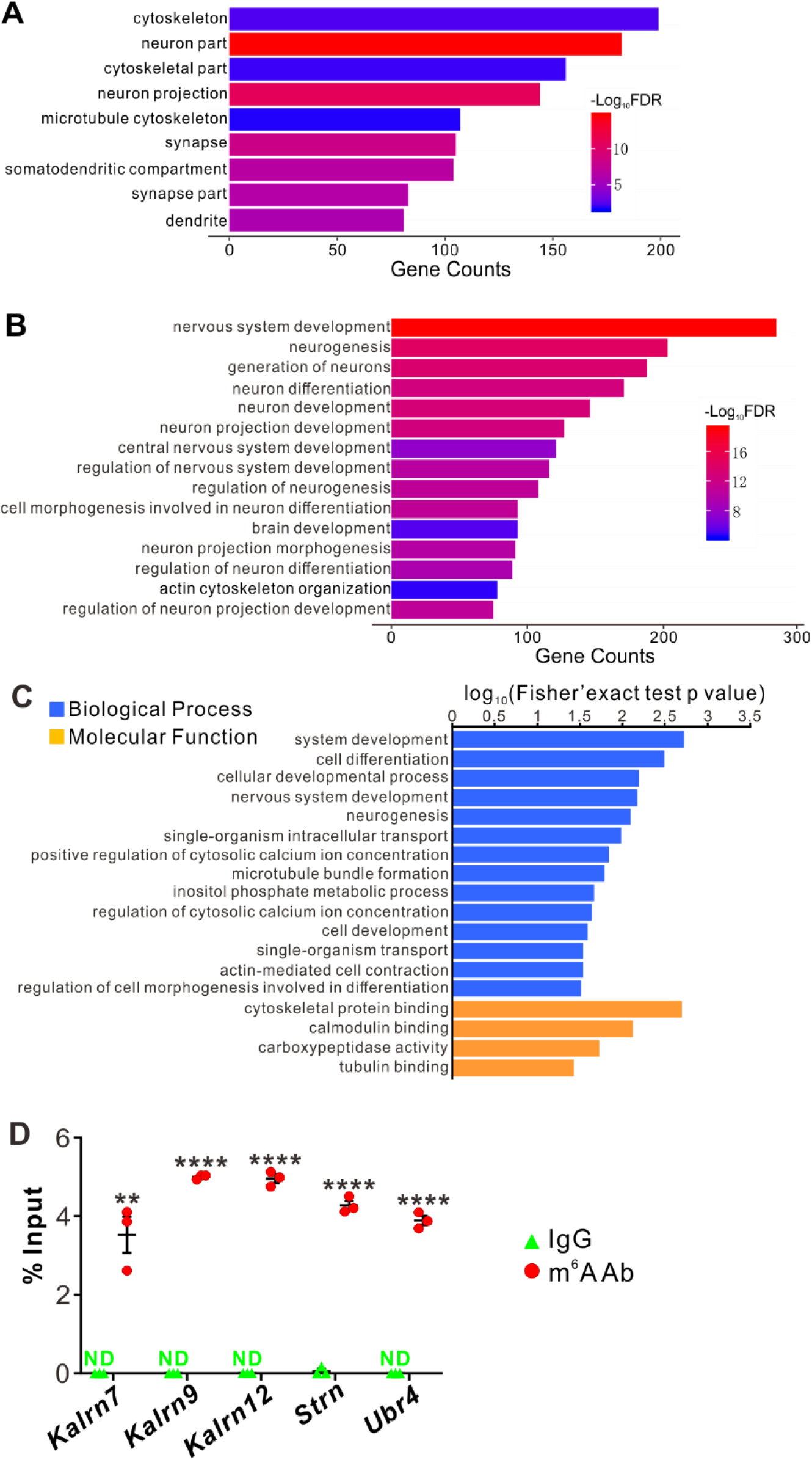
YTHDF2 target mRNAs were identified with transcriptomic and proteomic analysis. (**A, B**) Gene Ontology (GO) analysis of YTHDF2 target transcripts identified by anti-YTHDF2 RNA immunoprecipitation (RIP) in the retina followed by RNA sequencing (RIP-seq). Neural terms were picked out in Cellular Component (**A**) and Biological Process (**B**). (**C**) GO analysis of proteins which are upregulated after YTHDF2 KD by MS. (**D**) Verification of m^6^A modification of YTHDF2 target mRNAs by anti-m^6^A pulldown followed by RT-qPCR. ND, not detected. Data are mean ± SEM and are represented as dot plots (*n* = 3 replicates): \*\**p* = 0.0016 for *Kalrn7*; \*\*\*\**p* = 1.40E-08 for *Kalrn9*; \*\*\*\**p* = 1.46E-06 for *Kalrn12*; \*\*\*\**p* = 5.46E-06 for *Strn*; \*\*\*\**p* = 4.90E-06 for *Ubr4*; by unpaired Student’s *t* test.

The working model for YTHDF2 is that it binds and destabilizes its m^6^A-modified target transcripts ***(Wang et al. 2014)***. Since the destabilization of mRNAs will eventually decrease their protein levels, we carried out proteome analysis using mass spectrometry (MS) in acute *shYthdf2*-mediated knockdown of cultured RGCs, in order to identify directly affected targets. Three biological replicates of YTHDF2 knockdown (KD) followed by MS (YTHDF2 KD/MS) identified 114 proteins which were upregulated by YTHDF2 KD (*Supplementary file 2*). Functional annotation of these proteins revealed significant enrichment in neuron development- and cytoskeleton-related terms (*Figure 6C*), which is similar to anti-YTHDF2 RIP-Seq results.

By overlapping the two gene lists screened from anti-YTHDF2 RIP-Seq (*Supplementary file 1*) and YTHDF2 KD/MS_upregulation (*Supplementary file 2*), we identified a group of potential YTHDF2 target mRNAs in RGCs (*Supplementary file 3*), including *Kalrn*, *Strn* and *Ubr4*. m^6^A modification of these mRNAs were verified by anti m^6^A pull down (*Figure 6D*). *Kalrn* (*Kalirin*) gene generates three alternative splicing isoforms *Kalrn7*, *Kalrn9*, and *Kalrn12* encoding guanine-nucleotide exchange factors (GEFs) for Rho GTPases (Rho-GEF), which have been shown to regulate hippocampal and cortical dendritic branching ***(Xie et al. 2010; Yan et al. 2015)***, and are required for normal brain functions ***(Penzes et al. 2001; Xie et al. 2007; Cahill et al. 2009; Russell et al. 2014; Lu et al. 2015; Herring and Nicoll 2016)***. Strn (Striatin) was first identified in striatum, and functions as a B subunit of the serine/threonine phosphatase PP2A and is also a core component of a multiprotein complex called STRIPAK (striatin-interacting phosphatase and kinase complex) ***(Benoist et al. 2006; Li et al. 2018)***. Strn was reported to regulate dendritic arborization only in striatal neurons but not in cortical neurons ***(Li et al. 2018)***. However, whether and how Kalrn and Strn work in the retina was still unknown. Ubr4 (ubiquitin protein ligase E3 component N-recognin 4) is also known as p600 and has been shown to play roles in neurogenesis, neuronal migration, neuronal signaling and survival ***(Parsons et al. 2015)***. However, whether Ubr4 regulates dendrite development remains elusive.

### YTHDF2 controls the stability of its target mRNAs which encode proteins regulating RGC dendrite branching

MS analysis after YTHDF2 KD has shown that the protein levels of these target mRNAs were upregulated (*Supplementary file 2*). IF using antibodies against Strn and Ubr4 detected specific signals in the IPL which were increased in *Ythdf2* cKO retina compared with control retina (*Figure 7—figure supplement 1A*). Enrichment of these proteins in IPL implies that these proteins might function locally in RGC dendrites to regulate dendrite development.

We next wanted to know whether YTHDF2 controlled the protein levels of these m^6^A-modified target mRNAs through regulation of translation or transcript stability. As shown in *Figure 7—figure supplement 1B-D*, the mRNA levels of *Kalrn7*, *Kalrn9*, *Kalrn12*, *Strn* and *Ubr4* were dramatically increased after KD of YTHDF2 or METTL14, supporting that YTHDF2 might regulate stability of these target mRNAs. We further evaluated potential changes in the stability of these target mRNAs in an m^6^A-dependent manner. We further verified this by directly measuring the stability of these target mRNAs. As shown in *Figure 7A*, all the target mRNAs showed significantly increased stability in the *Ythdf2* cKO retina compared with controls. These results suggest that YTHDF2 controlled the protein levels of its m^6^A-modifed target mRNAs by decreasing their stability.

**Figure 7.**
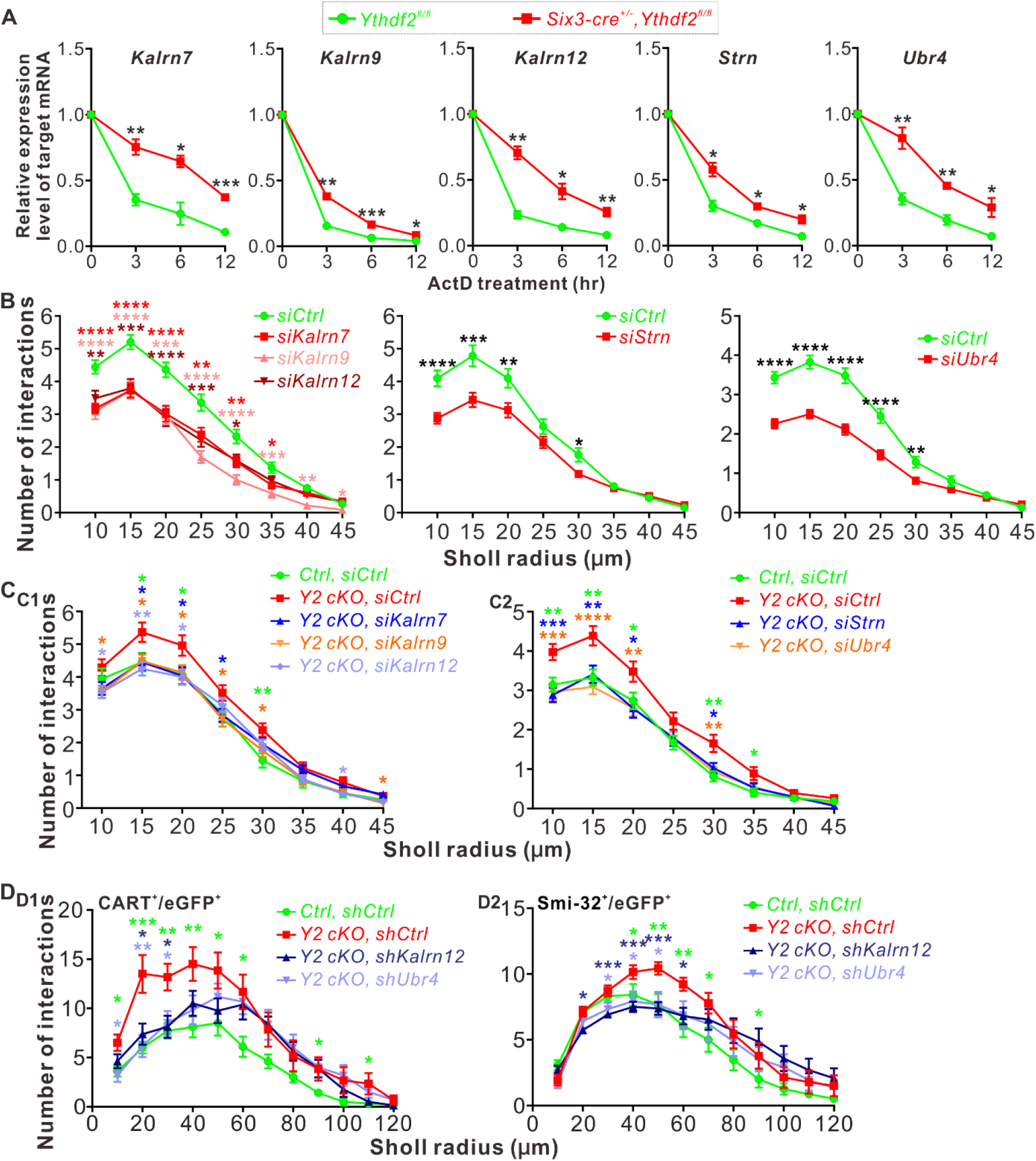
YTHDF2 target mRNAs mediate YTHDF2-controlled RGC dendrite branching. (**A**) YTHDF2 target mRNAs showing increased stability in the *Ythdf2* cKO retina. RGCs dissected from E14.5 *Ythdf2* cKO and control embryos were cultured, treated with actinomycin D (ActD) and collected at different timepoints. Data are mean ± SEM (*n* = 3 replicates). For *Kalrn7*, \**\*p* = 0.0057 (3 hr), *\*p* = 0.014 (6 hr), \*\**\*p* =0.00039 (12 hr); for *Kalrn9*, \**\*p* = 0.0036 (3 hr), \*\**\*p* = 0.00090 (6 hr), *\*p* =0.032 (12 hr); for *Kalrn12*, \**\*p* = 0.0012 (3 hr), *\*p* = 0.010 (6 hr), \**\*p* =0.0069 (12 hr); for *Strn*, *\*p* = 0.014 (3 hr), *\*p* = 0.012 (6 hr), *\*p* =0.016 (12 hr); for *Ubr4*, \**\*p* = 0.0077 (3 hr), \**\*p* = 0.0059 (6 hr), *\*p* =0.041 (12 hr); all by unpaired Student’s *t* test. (**B**) KD of the target mRNAs causing decreased dendrite branching of cultured RGCs prepared from wild type (WT) E14.5 retina by Sholl analysis. Brn3a and MAP2 IF were used to mark RGCs and visualize dendrites. Data are mean ± SEM. For *Kalrn7* (*n =* 59 for *siCtrl*, *n =* 56 for *siKalrn7*), \*\*\**\*p* = 2.33E-06 (10 μm), \*\*\**\*p* = 5.85E-06 (15 μm), \*\*\**\*p* = 8.67E-05 (20 μm), \**\*p* = 0.0045 (25 μm), \**\*p* = 0.0058 (30 μm), *\*p* = 0.010 (35 μm); for *Kalrn9* (*n =* 59 for *siCtrl*, *n =* 46 for *siKalrn9*), \*\*\**\*p* = 3.69E-05 (10 μm), \*\*\**\*p* = 5.53E-05 (15 μm), \*\**\*p* = 0.00020 (20 μm), \*\*\**\*p* = 3.09E-06 (25 μm), \*\*\**\*p* = 4.63E-06 (30 μm), \*\**\*p* = 0.00059 (35 μm), \**\*p* = 0.0010 (40 μm), *\*p* = 0.042 (45 μm); for *Kalrn12* (*n =* 59 for *siCtrl*, *n =* 39 for *siKalrn12*), \**\*p* = 0.0031 (10 μm), \*\**\*p* = 0.00017 (15 μm), \*\*\**\*p* = 6.56E-05 (20 μm), \**\*p* = 0.0017 (25 μm), *\*p* = 0.017 (30 μm); for *Strn* (*n =* 51 for *siCtrl*, *n =* 57 for *siStrn*), \*\*\**\*p* = 4.19E-05 (10 μm), \*\**\*p* = 0.00067 (15 μm), \**\*p* = 0.0079 (20 μm), *\*p* = 0.015 (30 μm); for *Ubr4* (*n =* 81 for *siCtrl*, *n =* 81 for *siUbr4*), \*\*\**\*p* = 1.26E-08 (10 μm), \*\*\**\*p* = 7.61E-10 (15 μm), \*\*\**\*p* = 2.35E-08 (20 μm), \*\*\**\*p* = 1.39E-05 (25 μm), \**\*p* = 0.0061 (30 μm); all by unpaired Student’s *t* test. (**C**) Increased dendrite branching of cultured RGCs prepared from E14.5 *Ythdf2* cKO retina was rescued by KD of target mRNAs using siRNAs. Data are mean ± SEM. *Ctrl*, *Ythdf2^fl/fl^*; *Y2 cKO*, *Six3-cre^+/-^, Ythdf2^fl/fl^*. In **C1**, “*Ctrl, siCtrl*” (*n* = 35 neurons) vs “*Y2 cKO, siCtrl*” (*n* = 52 neurons), \**p* = 0.038 (15 μm), \**p* = 0.045 (20 μm), \*\**p* = 0.0036 (30 μm); “*Y2 cKO, siKalrn7*” (*n* = 55 neurons) vs “*Y2 cKO, siCtrl*”, \**p* = 0.020 (15 μm), \**p* = 0.025 (20 μm), \**p* = 0.031 (25 μm); “*Y2 cKO, siKalrn9*” (*n* = 66 neurons) vs “*Y2 cKO, siCtrl*”, \**p* = 0.020 (10 μm), \**p* = 0.013 (15 μm), \**p* = 0.031 (20 μm), \**p* = 0.017 (25 μm), \**p* = 0.031 (30 μm), \**p* = 0.031 (45 μm); “*Y2 cKO, siKalrn12*” (*n* = 80 neurons) vs “*Y2 cKO, siCtrl*”, \**p* = 0.015 (10 μm), \*\**p* = 0.0018 (15 μm), \**p* = 0.015 (20 μm), \**p* = 0.027 (40 μm). In **C2**, “*Ctrl, siCtrl*” (*n* = 50 neurons) vs “*Y2 cKO, siCtrl*” (*n* = 47 neurons), \*\**p* = 0.0031 (10 μm), \*\**p* = 0.0013 (15 μm), \**p* = 0.029 (20 μm), \*\**p* = 0.0015 (30 μm), \**p* = 0.014 (35 μm); “*Y2 cKO, siStrn*” (*n* = 45 neurons) vs “*Y2 cKO, siCtrl*”, \*\*\**p* = 0.00016 (10 μm), \*\**p* = 0.0043 (15 μm), \**p* = 0.010 (20 μm), \**p* = 0.018 (30 μm); “*Y2 cKO, siUbr4*” (*n* = 57 neurons) vs “*Y2 cKO, siCtrl*”, \*\*\**p* = 0.00084 (10 μm), \*\*\*\**p* = 4.89E-05 (15 μm), \*\**p* = 0.0058 (20 μm), \*\**p* = 0.0045 (30 μm). All by unpaired Student’s *t* test. (**D**) Increased dendrite branching of RGC subtypes in *Ythdf2* cKO retina was rescued by KD of target mRNAs through intravitreal injection of AAV siRNAs in vivo. Data are mean ± SEM. *Ctrl*, *Ythdf2^fl/fl^*; *Y2 cKO*, *Six3-cre^+/-^,Ythdf2^fl/fl^*. In **D1** (CART^+^/eGFP^+^ ooDSGCs), “*Ctrl, shCtrl*” (*n* = 10 neurons) vs “*Y2 cKO, shCtrl*” (*n* = 6 neurons), \**p* = 0.010 (10 μm), \*\*\**p* = 0.00049 (20 μm), \*\**p* = 0.0021 (30 μm), \*\**p* = 0.0047 (40 μm), \**p* = 0.028 (50 μm), \**p* = 0.011 (60 μm), \**p* = 0.030 (90 μm), \**p* = 0.042 (110 μm); “*Y2 cKO, shKalrn12*” (*n* = 8 neurons) vs “*Y2 cKO, shCtrl*”, \**p* = 0.012 (20 μm), \**p* = 0.014 (30 μm); “*Y2 cKO, shUbr4*” (*n* = 6 neurons) vs “*Y2 cKO, shCtrl*”, \**p* = 0.011 (10 μm), \*\**p* = 0.0084 (20 μm), \**p* = 0.029 (30 μm). In **D2** (SMI-32^+^ αRGCs), “*Ctrl, shCtrl*” (*n* = 14 neurons) vs “*Y2 cKO, shCtrl*” (*n* = 14 neurons), \**p* = 0.032 (40 μm), \*\**p* = 0.0019 (50 μm), \*\**p* = 0.0014 (60 μm), \**p* = 0.015 (70 μm), \**p* = 0.044 (90 μm); “*Y2 cKO, shKalrn12*” (*n* = 26 neurons) vs “*Y2 cKO, shCtrl*”, \*\**p* = 0.0023 (20 μm), \*\*\**p* = 0.00076 (30 μm), \*\*\**p* = 0.00030 (40 μm), \*\*\**p* = 0.00020 (50 μm), \**p* = 0.015 (60 μm); “*Y2 cKO, shUbr4*” (*n* = 15 neurons) vs “*Y2 cKO, shCtrl*”, \**p* = 0.042 (30 μm), \**p* = 0.024 (40 μm), \**p* = 0.018 (50 μm). All by unpaired Student’s *t* test.

Next we explored the functions of these YTHDF2 target mRNAs in RGC dendrite development. We first generated siRNAs against these transcripts (*Figure 7—figure supplement 1E*). We then checked the effects on RGC dendrite branching after KD of these target mRNAs by siRNAs in cultured RGCs. As shown in *Figure 7B*, knockdown of *Kalrn7*, *Kalrn9*, *Kalrn12*, *Strn* or *Ubr4* led to significant decreases of RGC dendrite branching. Interestingly, the *siCocktail* against all these target mRNAs further significantly reduced the RGC dendrite branching compared with each individual siRNA (*Figure 7—figure supplement 1F*), suggesting that these targets may work in different pathways to regulate the RGC dendrite morphology. We further examined whether these target mRNAs mediate YTHDF2-regulated RGC dendrite branching. As shown in *Figure 2E-H*, and *Figure 3*, cKO of *Ythdf2* led to increased dendrite branching of RGCs both in vitro and in vivo. Transfection of siRNAs against these target mRNAs rescued dendrite branching increases in cultured *Ythdf2* cKO RGCs (*Figure 7C*). We continued to generate and performed intravitreal injection of AAV viral *shKalrn12* and *shUbr4*, which significantly rescued dendrite branching increases of CART^+^ ooDSGCs and SMI-32^+^ αRGCs in *Ythdf2* cKO retina in vivo (*Figure 7D*).

Taken together, we identified a group of YTHDF2 target mRNAs that encode proteins regulating RGC dendrite branching, which mediate YTHDF2-controlled RGC dendrite branching.

### *Ythdf2* cKO retina is more resistant to acute ocular hypertension (AOH)

The glaucomatous eyes are symptomatized with progressive neurodegeneration and vision loss ***(Agostinone and Di Polo 2015)***. High intraocular pressure is a major risk factor in glaucoma and has been shown to cause pathological changes in RGC dendrites before axon degeneration or soma loss is detected in different model animals ***(Weber et al. 1998; Shou et al. 2003; Morgan et al. 2006)***. Our findings that *Ythdf2* cKO in retina promotes RGC dendrite branching during development inspired us to wonder whether YTHDF2 also regulates RGC dendrite maintenance in the acute glaucoma model caused by acute ocular hypertension (AOH). We utilized the AOH model made with control and *Ythdf2* cKO mice to check whether *Ythdf2* cKO in the retina could alter the pathology in the glaucomatous eyes. RGC dendrite branching is significantly decreased after AOH operation compared with non-AOH in either genotype (*Figure 8—figure supplement 1A,B*). Interestingly, the *Ythdf2* cKO retina with AOH operation maintains significantly higher dendrite complexity compared with the glaucomatous eyes of *Ythdf2^fl/fl^* control mice (*Figure 8A,B*). In addition, there are significant RGC neuron losses in both genotypes after AOH (*Figure 8C,D*). However, the reduction of RGC number in the *Ythdf2* cKO retina is less than control retina (*Figure 8C,D*). These results support that *Ythdf2* cKO protects retina from RGC dendrite degeneration and soma loss caused by AOH.

**Figure 8.**
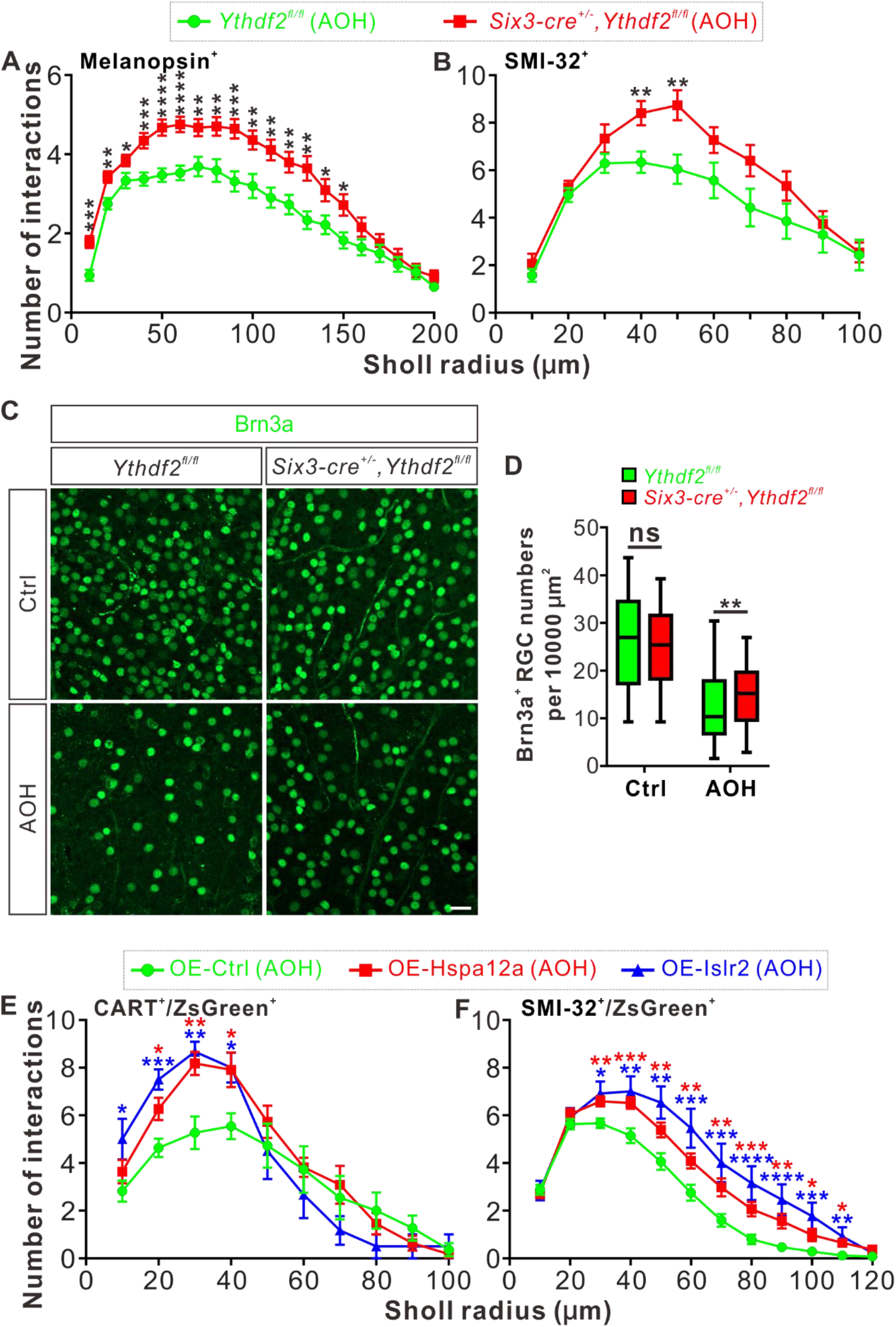
*Ythdf2* cKO retina is more resistant to acute ocular hypertension (AOH). (**A, B**) Better maintenance of RGC dendrite arborization in *Ythdf2* cKO retina after AOH operation. AOH was performed using adult mice, and retinas were collected after AOH for wholemount immunostaining of melanopsin and SMI-32 to visualize the dendrite arbors of corresponding RGC subtype, respectively. Dendrite traces were drawn as previously shown and quantification of dendrite branching was done using Sholl analysis. Data are mean ± SEM. Numbers of interactions are significantly greater in *Six3-cre^+/-^,Ythdf2^fl/fl^* retina than *Ythdf2^fl/fl^* control retina in both RGC subtypes after AOH: for melanopsin^+^ ipRGCs in **A**, *Ythdf2^fl/fl^*/AOH (*n* = 51 RGCs) vs cKO/AOH (*n* = 64 RGCs), \*\*\**p* = 0.00015 (10 μm), \*\**p* = 0.0017 (20 μm), \**p* = 0.034 (30 μm), \*\*\**p* = 0.00035 (40 μm), \*\*\*\**p* = 3.02E-05 (50 μm), \*\*\*\**p* = 2.63E-05 (60 μm), \*\**p* = 0.0029 (70 μm), \*\**p* = 0.0028 (80 μm), \*\*\**p* = 0.00035 (90 μm), \*\**p* = 0.0032 (100 μm), \*\**p* = 0.0014 (110 μm), \*\**p* = 0.0043 (120 μm), \*\**p* = 0.0014 (130 μm), \**p* = 0.023 (140 μm), \**p* = 0.013 (150 μm); for SMI-32^+^ αRGCs in **B**, *Ythdf2^fl/fl^*/AOH (*n* = 21 neurons) vs cKO/AOH (*n* = 15 neurons), \*\**p* = 0.0052 (40 μm), \*\**p* = 0.0057 (50 μm); all by unpaired Student’s *t* test. (**C, D**) *Ythdf2* cKO retina showing less severe RGC loss after AOH. AOH was performed using adult mice and retinas were collected after AOH for wholemount immunostaining using a Brn3a antibody (**C**). Numbers of Brn3a^+^ RGCs per 10000 μm^2^ of retina were quantified for different genotypes and conditions (confocal fields for analysis: *n =* 117 for *Ythdf2^fl/fl^*/Ctrl; *n =* 98 for *Ythdf2^fl/fl^*/AOH; *n =* 110 for cKO/Ctrl; *n =* 104 for cKO/AOH). Data are represented as box and whisker plots (**D**): ns, not significant (*p* = 0.16; *Ythdf2^fl/fl^*/Ctrl vs cKO/Ctrl); \**\*p* = 0.0077 (*Ythdf2^fl/fl^*/AOH vs cKO/AOH); by unpaired Student’s *t* test. Scale bar: 25 μm. (**E, F**) Overexpression (OE) of YTHDF2 targets *Hspa12a* and *Islr2* protecting retina from RGC dendrite degeneration. Wild type (WT) mice were intravitreally injected with AAV overexpressing *Hspa12a* or *Islr2* and then operated with AOH. Wholemount immunostaining of CART/ZsGreen and SMI-32/ZsGreen was carried out to visualize the dendrite arbors of corresponding RGC subtype, respectively. Dendrite traces were drawn as previously shown and quantification of dendrite branching was done using Sholl analysis. Data are mean ± SEM. Numbers of interactions are significantly greater in retina with OE of *Hspa12a* or *Islr2* than control retina in both RGC subtypes after AOH. For CART^+^ ooDSGCs in **E**: OE-Ctrl/AOH (*n* = 11 RGCs) vs OE-Hspa12a/AOH (*n* = 11 RGCs), *p = 0.014 (20 μm), \*\**p* = 0.0025 (30 μm), \**p* = 0.018 (40 μm); OE-Ctrl/AOH vs OE-Islr2/AOH (*n* = 6 RGCs), \**p* = 0.024 (10 μm), \*\*\**p* = 0.00031 (20 μm), \*\**p* = 0.0038 (30 μm), \**p* = 0.013 (40 μm). For SMI-32^+^ αRGCs in **F**: OE-Ctrl/AOH (*n* = 49 neurons) vs OE-Hspa12a/AOH (*n* = 46 neurons), \*\**p* = 0.0023 (30 μm), \*\*\**p* = 0.00080 (40 μm), \*\**p* = 0.0059 (50 μm), \*\**p* = 0.0051 (60 μm), \*\**p* = 0.0036 (70 μm), \*\*\**p* = 0.00070 (80 μm), \*\**p* = 0.0015 (90 μm), \**p* = 0.016 (100 μm), \**p* = 0.011 (110 μm); OE-Ctrl/AOH vs OE-Islr2/AOH (*n* = 13 RGCs), \**p* = 0.010 (30 μm), \*\**p* = 0.0093 (40 μm), \*\**p* = 0.0019 (50 μm), \*\*\**p* = 0.00085 (60 μm), \*\*\**p* = 0.00067 (70 μm), \*\*\*\**p* = 4.25E-05 (80 μm), \*\*\*\**p* = 2.54E-05 (90 μm), \*\*\**p* = 0.00020 (100 μm), \*\**p* = 0.0016 (110 μm). All by unpaired Student’s *t* test.

Next we wanted to know whether and how YTHDF2 target mRNAs mediate these effects in the AOH models. We first checked the expression of YTHDF2 target mRNAs identified in the developing retina (*Supplementary file 3*) in the adult *Ythdf2* cKO and control retina. We found that two target mRNAs *Hspa12a* and *Islr2* show upregulation in the adult *Ythdf2* cKO retina compared with control (*Figure 8—figure supplement 1C*). m^6^A modification of *Hspa12a* and *Islr2* mRNAs was further verified by anti-m^6^A pulldown (*Figure 8—figure supplement 1D*). *Hspa12a* encodes heat shock protein A12A which is an atypical member of the heat shock protein 70 family and has been shown to be downregulated in diseases such as ischemic stroke, schizophrenia, and renal cell carcinoma ***(Pongrac et al. 2004; Mao et al. 2018; Min et al. 2020)***. *Islr2* encodes immunoglobulin superfamily containing leucine-rich repeat protein 2 and is poorly studied. Here, we found that *Hspa12a* and *Islr2* are downregulated in the retina after AOH operation (*Figure 8—figure supplement 1E*), which is likely caused by upregulation of YTHDF2 in the AOH-treated retina (*Figure 8—figure supplement 1F-H*). We therefore hypothesized that AOH upregulates YTHDF2 which in turn downregulates its targets *Hspa12a* and *Islr2*, thus causing RGC dendrite degeneration and soma loss. If this is the case, overexpression of *Hspa12a* and *Islr2* might protect RGC dendrite from AOH-triggered degeneration. We thus generated AAV harboring overexpression constructs of *Hspa12a* and *Islr2* which were intravitreally injected to wild type retinas. After the AOH induction, the retinas overexpressing *Hspa12a* and *Islr2* maintain significantly more complex RGC dendrite arbor compared with control AAV (*Figure 8E,F*).

These data verify that loss-of-function of YTHDF2 and gain-of-function of its targets *Hspa12a* and *Islr2* have neuroprotective roles in the glaucomatous retina.

## Discussion

Functions and mechanisms of mRNA m^6^A modification in the dendrite development were not known. Here, we revealed a critical role of the m^6^A reader YTHDF2 in RGC dendrite development and maintenance. YTHDF2 have two phases of function to control RGC dendrite development first and then maintenance through regulating two sets of target mRNAs. In early postnatal stages, the target mRNAs *Kalrn7*, *Kalrn9*, *Kalrn12*, *Strn* and *Ubr4* mediate YTHDF2 functions to regulate RGC dendrite development. In the adult mice, another set of target mRNAs *Hspa12a* and *Islr2* mediate YTHDF2 function to regulate RGC dendrite maintenance.

### Positive and negative regulators for dendrite development

The general principle for dendrite arborization is that the dendrite arbor cannot be either too big or too small in order to precisely sample a presynaptic target area during neural circuit formation ***(Lefebvre et al. 2015)***. Numerous extrinsic and intrinsic mechanisms have been found to regulate dendritic arbor patterning, which involves both positive and negative factors to achieve balanced control of dendritic growth ***(Jan and Jan 2010; Dong et al. 2015; Ledda and Paratcha 2017)***. For the secreted and diffusible cues, BDNF promotes dendrite branching and complexity ***(Cheung et al. 2007)***; the non-canonical Wnt7b/PCP pathway is a positive regulator of dendrite growth and branching ***(Rosso et al. 2005)***; the non-canonical Wnt receptor Ryk works as a negative regulator by limiting the extent of dendritic branching ***(Lanoue et al. 2017)***. For the contact-mediated signals, the cadherins Celsr2 and Celsr3 regulate dendrite growth in an opposite manner in cortical pyramidal and Purkinje neurons, and hippocampal neurons, respectively ***(Shima et al. 2004; Shima et al. 2007)***. For the transcription factors, studies have shown that manipulation of Cux1 and Cux2 levels has distinct effects on apical and basal arbors of cortical dendrites ***(Cubelos et al. 2015)***; interestingly, the functions of Sp4 in dendrite development are dependent on the cellular context of its expression, e.g. Sp4 promotes dendrite growth and branching in hippocampal dentate granule cells but limits dendrite branching in cerebellar granule cells ***(Ramos et al. 2007; Zhou et al. 2007)***. Here we identified another negative regulator YTHDF2 which works posttranscriptionally, and loss-of-function of YTHDF2 increased dendrite complexity during development and protected RGC degeneration from AOH.

### Posttranscriptional regulation of dendrite development

It is well established that mRNAs can be transported and targeted to specific neuronal compartments such as axons and dendrites. Local translation of these mRNAs enables exquisite and rapid control of local proteome in specific subcellular compartments ***(Ledda and Paratcha 2017)***. Local translation is known to play roles in controlling dendrite arborization ***(Chihara et al. 2007)***, and is regulated by specific RNA-binding proteins ***(Jan and Jan 2010)***. In Drosophila, the RNA binding proteins Pumilio (Pum), Nanos (Nos), Glorund (Glo) and Smaug (Smg) regulate morphogenesis and branching of specific classes of dendritic arborization (da) neurons through controlling translation of their target mRNAs including *nanos* mRNA itself ***(Ye et al. 2004; Brechbiel and Gavis 2008)***. The mouse homologue of another RNA-binding protein Staufen, Stau1, regulates dendritic targeting of ribonucleoprotein particles and dendrite branching ***(Vessey et al. 2008)***. Here we found that the m^6^A reader and RNA-binding protein YTHDF2 controls stability of its target mRNAs and regulates dendrite branching in RGCs. It would be interesting to see whether these target mRNAs are localized into dendrites and whether YTHDF2 works in dendrites to control their stability and translation. Actually, *Strn4* mRNA has been shown to be present in dendrites and locally translated ***(Lin et al. 2017)***. In addition, how the proteins encoded by these target mRNAs regulate RGC dendrite branching during development and maintenance remains to be explored and will be important future directions.

### Neuroprotective genes in retinal injuries and degeneration

Transcriptome analyses have revealed differentially expressed genes after retinal injuries such as AOH-induced glaucoma and optic nerve crush (ONC), and the up-regulated genes are of importance for discovering new treatment approaches ***(Jakobs 2014; Tran et al. 2019)***. One of the previous studies has identified *Mettl3*, encoding the m^6^A writer, as an up-regulated gene after ONC ***(Agudo et al. 2008)***. Here we found *Ythdf2*, encoding an m^6^A reader, was also up-regulated in the retina after AOH. We further found that *Hspa12a* and *Islr2*, two targets of YTHDF2 in adult retina, were downregulated in glaucomatous retinas. Overexpression of *Hspa12a* and *Islr2* protected retina from AOH-caused RGC dendrite degeneration. Our findings in this study suggest that YTHDF2 and its neuroprotective target mRNAs might be valuable in developing novel therapeutic approaches to treat neurodegeneration caused by glaucoma and other retinal injuries.

## Materials and methods

### Key resources table

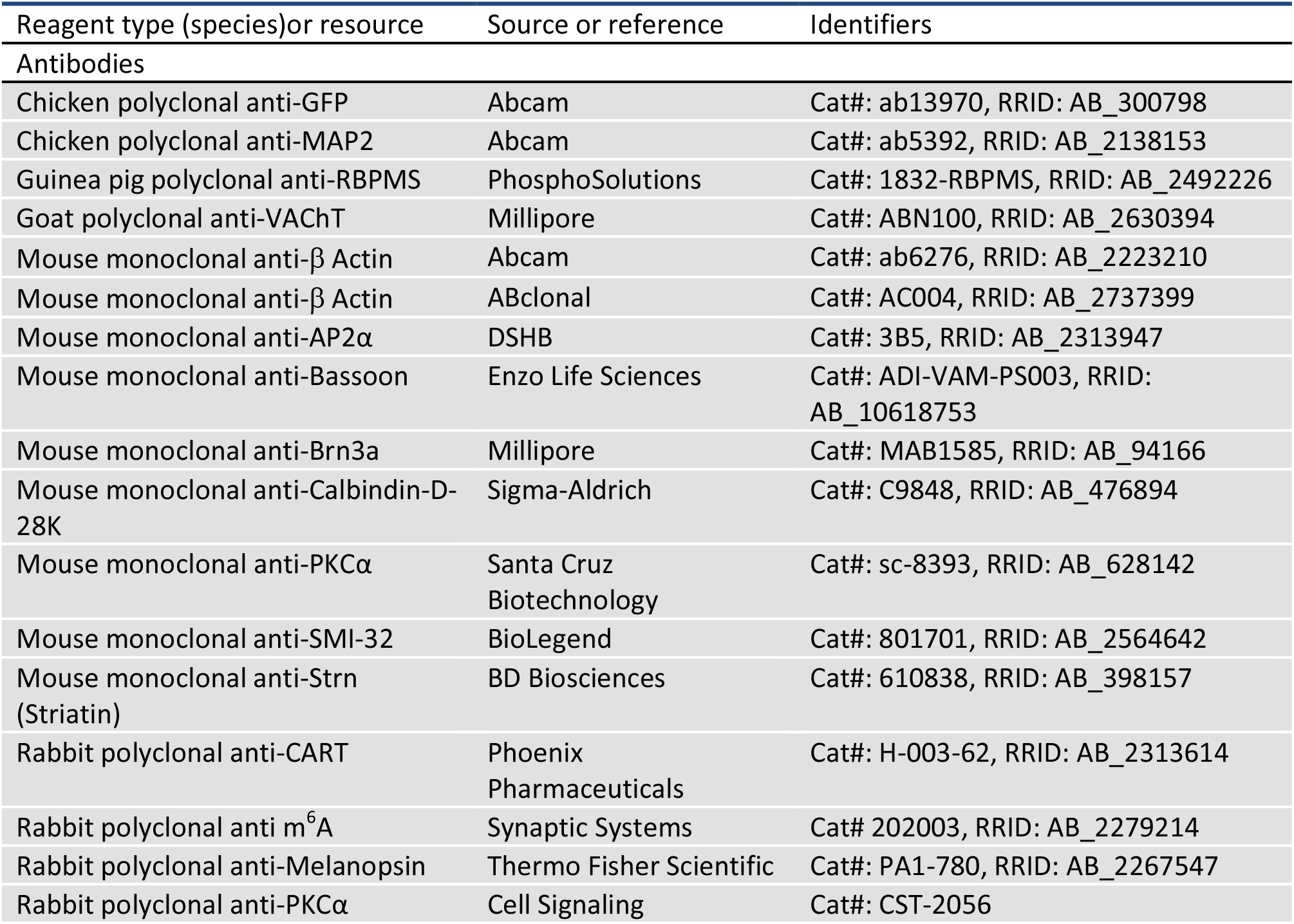
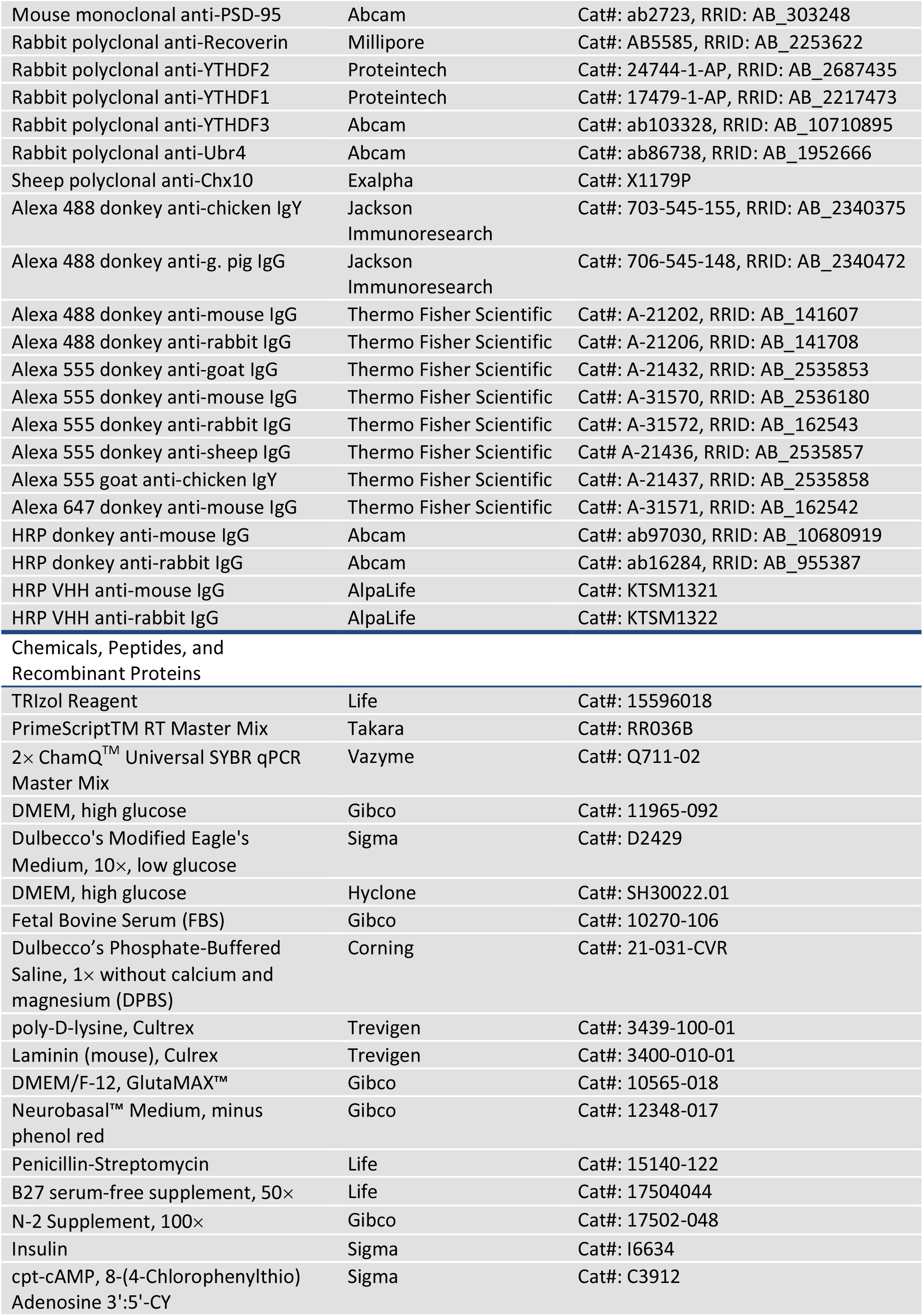
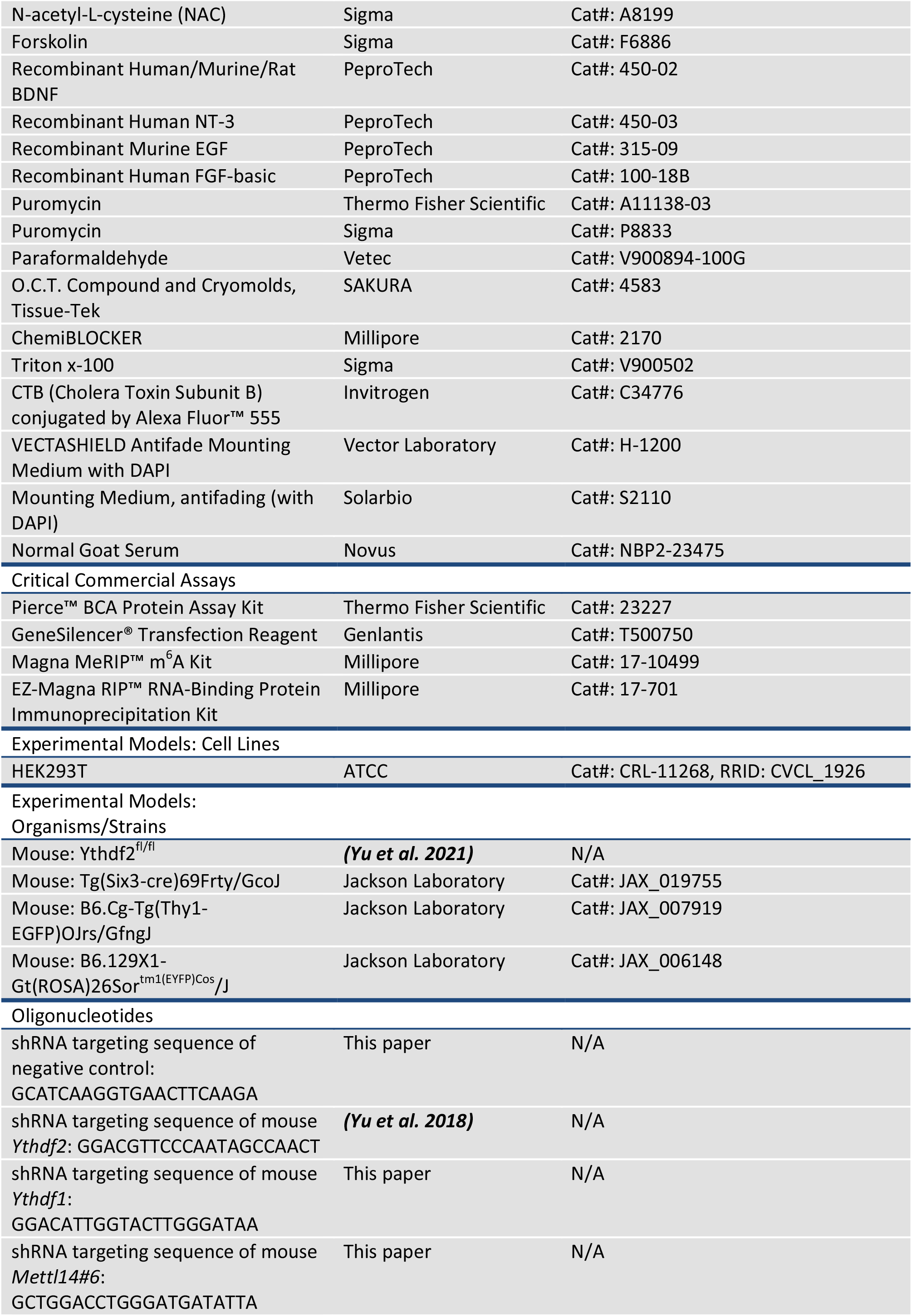
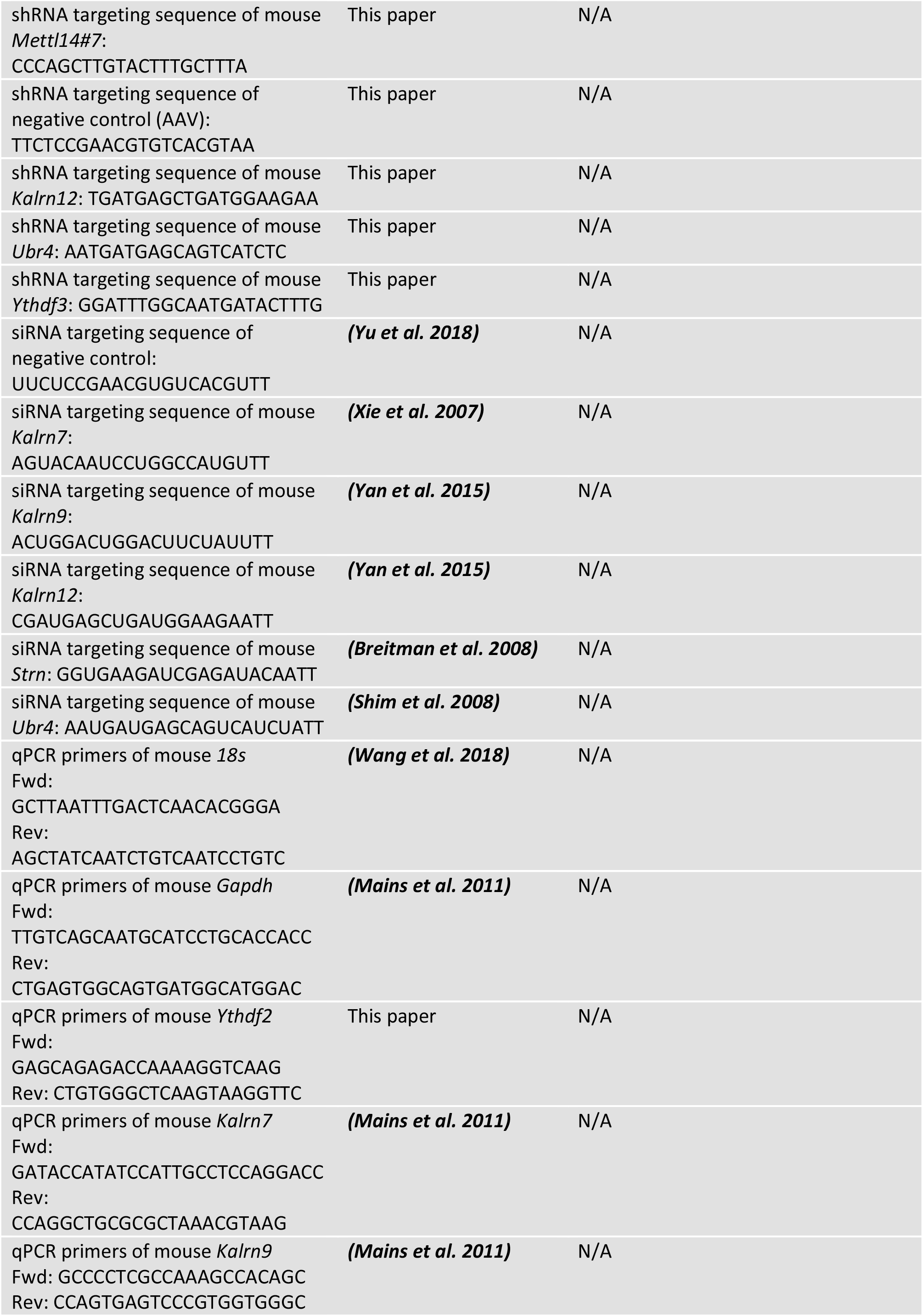
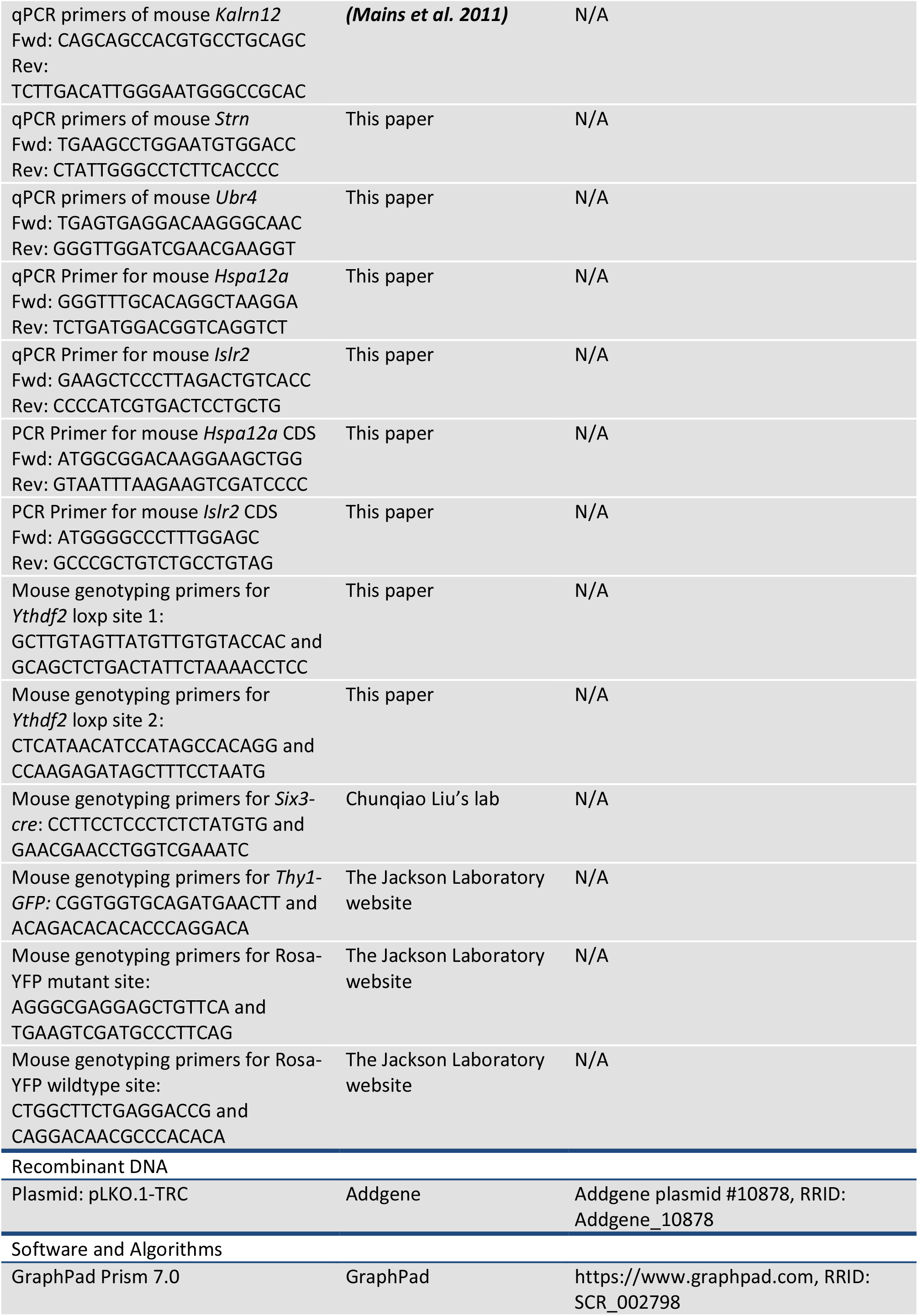
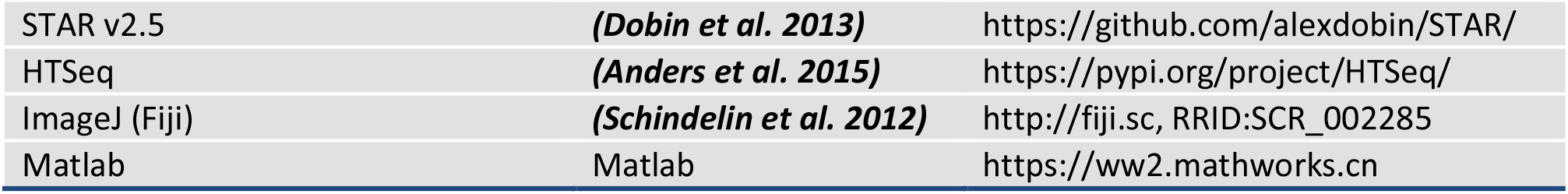

### Animals and generation of the *Ythdf2* cKO mice

*Ythdf2^fl/fl^* mice were reported previously ***(Yu et al. 2021)***. *Six3-cre **(Furuta et al. 2000)***, *Thy1-GFP **(Feng et al. 2000)*** and *Rosa26-eYFP **(Srinivas et al. 2001)*** mice were from Jackson Laboratory. For timed pregnancy, embryos were identified as E0.5 when a copulatory plug was observed. Genotyping primers are as following: the first *Ythdf2-loxP* site, 5’-GCTTGTAGTTATGTTGTGTACCAC-3’ and 5’-GCAGCTCTGACTATTCTAAAACCTCC-3’; the second *Ythdf2-loxP* site, 5’-CTCATAACATCCATAGCCACAGG-3’ and 5’-CCAAGAGATAGCTTTCCTAATG-3’. *Six3-cre* site, 5’-CCTTCCTCCCTCTCTATGTG-3’ and 5’-GAACGAACCTGGTCGAAATC-3’. *Rosa26-eYFP* wild type site, 5’-CTGGCTTCTGAGGACCG-3’ and 5’-CAGGACAACGCCCACACA-3’; the mutant site, 5’-AGGGCGAGGAGCTGTTCA-3’ and 5’-TGAAGTCGATGCCCTTCAG-3’. All experiments using mice were carried out following the animal protocols approved by the Laboratory Animal Welfare and Ethics Committee of Southern University of Science and Technology.

### Retinal neuronal culture

Retinal neurons were dissociated from E14.5-15.5 mouse embryos by papain in DPBS (1× Dulbecco’s Phosphate-Buffered Saline, Corning) following the previously described methods ***(Kechad et al. 2012)***, and neuronal suspension was plated on acid-washed glass coverslips pre-coated with poly-D-lysine (Trevigen, 100 μg/ml) for 1 hr and laminin (Trevigen, 5 μg/ml) overnight at 37°C. Culture medium was made up of half DMEM/F12 medium (Gibco) and half neurobasal medium (Gibco), supplemented with B27 supplement (Life, 0.5×), penicillin-streptomycin (Life, 1×), N-2 supplement (Gibco, 0.5×), N-acetyl-L-cysteine (Sigma, NAC 0.6 mg/ml), cpt-cAMP (Sigma, 100 μM), forskolin (Sigma, 10 μM), and insulin (Sigma, 25 μg/ml). EGF (PeproTech, 50 ng/ml), BDNF (PeproTech, 50 ng/ml), NT-3 (PeproTech, 25 ng/ml), and FGF-basic (PeproTech, 10 ng/ml) were freshly added before using.

### Knockdown using lentiviral shRNA, siRNA or AAV shRNA, and overexpression using AAV system

Lentiviral knockdown plasmids encoding shRNA (*shCtrl*: 5’-GCATCAAGGTGAACTTCAAGA-3’; *shYthdf2*: 5’-GGACGTTCCCAATAGCCAACT-3’; *shYthdf1:* 5’-GGACATTGGTACTTGGGATAA-3’; *shYthdf3*: 5’-GGATTTGGCAATGATACTTTG-3’; *shMettl14#6:* 5’-GCTGGACCTGGGATGATATTA-3’; *shMettl14#7:* 5’-CCCAGCTTGTACTTTGCTTTA-3’) were generated from pLKO.1-TRC and lentivirus preparation process was described previously ***(Yu et al. 2018)***. All siRNA were chosen from previous studies and the target sequences of siRNA are as following: *siCtrl* (RNAi negative control): 5’-UUCUCCGAACGUGUCACGUTT-3’ ***(Yu et al. 2018)***; *siKalrn7*: 5’-AGUACAAUCCUGGCCAUGUTT-3’ ***(Xie et al. 2007)***; *siKalrn9*: 5’-ACUGGACUGGACUUCUAUUTT-3’ ***(Yan et al. 2015)***; *siKalrn12*: 5’-CGAUGAGCUGAUGGAAGAATT-3’ ***(Yan et al. 2015)***; *siStrn*: 5’-GGUGAAGAUCGAGAUACAATT-3’ ***(Breitman et al. 2008)***; *siUbr4*: 5’-AAUGAUGAGCAGUCAUCUATT-3’ ***(Shim et al. 2008)***. AAV knockdown plasmids encoding shRNA (*shCtrl*: 5’-TTCTCCGAACGTGTCACGTAA-3’; *shKalrn12*: 5’-TGATGAGCTGATGGAAGAA-3’; *shUbr4*: 5’-AATGATGAGCAGTCATCTC-3’) were generated using pHBAAV-U6-MCS-CMV-EGFP and packaged in serotype-9 by Hanbio (1.5×10^12^ genomic copies per ml). AAV overexpression plasmids of Hspa12a (NM_175199.3; PCR primer for mouse *Hspa12a*: 5’-ATGGCGGACAAGGAAGCTGG-3’ and 5’-GTAATTTAAGAAGTCGATCCCC-3’) and Islr2 (NM_001161541.1; PCR Primer for mouse *Islr2*: 5’-ATGGGGCCCTTTGGAGC-3’ and 5’-GCCCGCTGTCTGCCTGTAG-3’) were generated from pHBAAV-CMV-MCS-3flag-T2A-ZsGreen and packaged serotype-9 by Hanbio (1.2×10^12^ genomic copies per ml).

GeneSilencer® Transfection Reagent (Genlantis) was used in siRNA transfection following the manufacturer’s protocols. Culture medium was changed after 1 day of lentiviral shRNA infection or siRNA transfection. For lentiviral shRNA assay, puromycin (Thermo or Sigma, 1 μg/ml) was added after 2 days of infection. Immunofluorescence, RNA or protein preparation was performed after shRNA or siRNA worked for 3 days. For AAV intravitreal injection, P0-P1 mouse pups were anesthetized in ice and then eyes were pierced at the edge of corneal by 30G × 1/2 needle (BD, 305106) under stereomicroscope. Then 1 μl AAV was intravitreally injected with 10 µL Syringe (Hamilton, 80330) following the pinhole. P15 or adult mice were anesthetized with 2.5% Avertin and then eyes were pierced at the side of corneal and the outer segment of sclera by 30G × 1/2 needle successively. 2 μl AAV was intravitreally injected with 10 µL Syringe following the pinhole on the sclera. All subsequent experiments such as acute ocular hypertension operation and immunostaining were carried out after at least 3 weeks (10 days for ZsGreen/CART labeling of ooDSGCs in *Ythdf2* cKO and control mice in *Figure 3A*).

### RT-qPCR

Total RNA was extracted from cells or tissues with TRIzol Reagent (Life) and then used for reverse transcription by PrimeScript^TM^ RT Master Mix (TaKaRa). Synthesized cDNA was used for qPCR by 2× ChamQ^TM^ Universal SYBR qPCR Master Mix (Vazyme) on StepOnePlus^TM^ Real-Time PCR System (ABI) or BioRad CFX96 Touch Real-Time PCR system. Primers used for qPCR are as following: mouse *Gapdh*: 5’-TTGTCAGCAATGCATCCTGCACCACC-3’ and 5’-CTGAGTGGCAGTGATGGCATGGAC-3’ ***(Mains et al. 2011)***; mouse *Kalrn7*: 5’-GATACCATATCCATTGCCTCCAGGACC-3’ and 5’-CCAGGCTGCGCGCTAAACGTAAG-3’ ***(Mains et al. 2011)***; mouse *Kalrn9*: 5’-GCCCCTCGCCAAAGCCACAGC-3’ and 5’-CCAGTGAGTCCCGTGGTGGGC-3’ ***(Mains et al. 2011)***; mouse *Kalrn12*: 5’-CAGCAGCCACGTGCCTGCAGC-3’ and 5’-TCTTGACATTGGGAATGGGCCGCAC-3’ ***(Mains et al. 2011)***; mouse *Strn*: 5’-TGAAGCCTGGAATGTGGACC-3’ and 5’-CTATTGGGCCTCTTCACCCC-3’; mouse *Ubr4*: 5’-TGAGTGAGGACAAGGGCAAC-3’ and 5’-GGGTTGGATCGAACGAAGGT-3’; mouse *Ythdf2*: 5’-GAGCAGAGACCAAAAGGTCAAG-3’and 5’-CTGTGGGCTCAAGTAAGGTTC-3’; 18s: 5’-GCTTAATTTGACTCAACACGGGA-3’ and 5’-AGCTATCAATCTGTCAATCCTGTC-3’ ***(Wang et al. 2018)***; mouse *Hspa12a*: 5’-GGGTTTGCACAGGCTAAGGA-3’ and 5’-TCTGATGGACGGTCAGGTCT-3’; mouse *Islr2*: 5’-GAAGCTCCCTTAGACTGTCACC-3’ and 5’-CCCCATCGTGACTCCTGCTG-3’.

### Immunofluorescence and immunostaining

For tissue sections, mouse embryonic eyes were fixed with 4% PFA (Sigma) in 0.1 M Phosphate Buffer (PB) for 30-45 min at room temperature (RT); eyes of mouse pups (< P10) were pre-fixed briefly and then eyecups were dissected and fixed for 45 min-1 hr at RT; for P20-30 or adult mice, eyecups were dissected after myocardial perfusion with 0.9% NaCl, followed by fixation for 1 hr. After PBS (3 × 5 min) washing, tissues were dehydrated with 30% sucrose in 0.1 M PB overnight at 4°C, then embedded with O.C.T. (SAKURA) and cryosectioned at 12 μm (20 μm for Thy1-GFP section analysis) with Leica CM1950 Cryostat. Tissue sections were permeabilized and blocked with 10% ChemiBLOCKER (Millipore) and 0.5% Triton x-100 (Sigma) in PBS (PBST) for 1 hr at RT and incubated in PBST overnight at 4°C with following primary antibodies: chicken anti-GFP (1:1000, Abcam ab13970), chicken anti-MAP2 (1:10000, Abcam ab5392), goat anti-VAChT (1:1000,Millipore ABN100), guinea pig anti-RBPMS (1:1000, PhosphoSolutions 1832-RBPMS), mouse anti-AP2α (1:1000, DSHB 3B5), mouse anti-Bassoon (1:2500, Enzo Life Sciences ADI-VAM-PS003), mouse anti-Brn3a (1:300, Millipore MAB1585), mouse anti-Calbindin-D-28K (1:200, Sigma C9848), mouse anti-PKCα (1:500, Santa Cruz sc-8393), rabbit anti-Strn (Striatin) (1:500, BD Biosciences 610838), rabbit anti-CART (1:2000, Phoenix Pharmaceuticals H-003-62), rabbit anti-m^6^A (1:200, Synaptic Systems 202003), rabbit anti-melanopsin (1:1000, Thermo PA1-780), rabbit anti-PKCα (1:1000, Cell Signaling CST-2056), rabbit anti-PSD95 (1:1000, Abcam ab18258), rabbit anti-Recoverin (1:1000, Millipore AB5585), rabbit anti-YTHDF2 (1:1000, Proteintech 24744-1-AP), rabbit anti-YTHDF1 (1:1000, Proteintech 17479-1-AP), rabbit anti-YTHDF3 (1:1000, Abcam ab103328), rabbit anti-Ubr4 (1:300, Abcam ab86738), sheep anti-Chx10 (1:1000, Exalpha X1179P). After three times of PBS washing, sections were incubated in PBST for 1 hr at RT with secondary antibodies: Alexa 488 donkey anti-chicken (1:500, Jackson 703-545-155), Alexa 488 donkey anti guinea pig (1:500, Jackson 706-545-148), Alexa 488 donkey anti-mouse (1:500, Thermo A21202), Alexa 488 donkey anti-rabbit (1:500, Thermo A21206), Alexa 555 donkey anti-goat (1:1000, Thermo A21432), Alexa 555 donkey anti-mouse (1:1000, Thermo A31570), Alexa 555 donkey anti-rabbit (1:1000, Thermo A31572), Alexa 555 donkey anti-sheep (1:1000, Thermo A21436), Alexa 555 goat anti-chicken (1:1000, Thermo A21437), or Alexa 647 donkey anti-mouse (1:200, Thermo A31571) and then mounted with the VECTASHIELD Antifade Mounting Medium with DAPI (Vector Laboratory).

For cultured neurons, after twice of PBS washing, cells were fixed for 15 min with 4% PFA in 0.1 M PB at RT, then washed with PBS three times and blocked in PBST for 20 min at RT. Antibody incubation conditions are the same as tissue sections.

For wholemount immunostaining of retina, eyes were dissected after myocardial perfusion with 0.9% NaCl. Then retinas were separated from sclera and fixed with 4% PFA in 0.1M PB for 1 hr at RT. Then retinas were blocked with 5% normal goat serum (Novus), 0.4% Triton x-100 in PBS overnight at 4°C. Primary antibodies such as chicken anti-GFP (1:1000, Abcam ab13970), mouse anti-Brn3a (1:300, Millipore MAB1585), mouse anti-SMI-32 (1:200, BioLegend 801701), or rabbit anti-Melanopsin (1:1000, Thermo PA1-780), rabbit anti-CART (1:2000, Phoenix Pharmaceuticals H-003-62) were diluted in 5% normal goat serum, 0.4% Triton x-100 in PBS and incubated overnight at 4°C. Then retinas were incubated with Alexa 488 donkey anti-chicken (1:500, Jackson 703-545-155), Alexa 488 donkey anti-mouse (1:500, Thermo A21202), Alexa 555 donkey anti-mouse (1:1000, Thermo A-31570) and Alexa 555 donkey anti-rabbit (1:1000, Thermo A31572) second antibodies in 5% normal goat serum (Novus), 0.4% Triton x-100 in PBS and finally mounted with the VECTASHIELD Antifade Mounting Medium with DAPI.

All images were captured on Nikon A1R confocal microscope or Zeiss LSM 800 confocal microscope with identical settings for each group in the same experiment. A region of interest (ROI), length or thickness in immunofluorescence experiments were obtained with ImageJ. The number of neurons in specific area was counted blindly and manually. To quantify RGC dendrite lamination in IPL with Thy1-GFP, z-stack and maximum projection were performed during the analysis. GFP intensity values across IPL depth were measured by ImageJ/Analyze/Plot Profile function ***(Liu et al. 2018)***. To quantify the numbers of Bassoon^+^/PSD-95^+^ excitatory synapses in IPL, the colocalization puncta was measured by ImageJ/Analyze/Puncta Analyzer as described previously ***(Ippolito and Eroglu 2010)***.

### Sholl analysis

For confocal images of cultured RGCs, MAP2 signals in original format were analyzed with simple neurite tracer and then quantified with Sholl analysis (5 μm per distance from soma center) which was a widely used method in neurobiology to quantify the complexity of dendritic arbors using ImageJ ***(Schindelin et al. 2012; Binley et al. 2014)***. Retina wholemount data were captured in z-stack mode (0.5-1 μm per slide) with confocal microscopes. ZsGreen, eGFP and SMI-32 signals were directly analyzed with simple neurite tracer and then z projection of all tracers was quantified with Sholl analysis (10 μm per distance from soma center), while melanopsin signals were maximum-projected before tracing.

### Optomotor response assay

*Ythdf2* cKO and control mice aged about 6 weeks were dark-adapted overnight before experiment and used in the optomotor response assay following the previously reported protocols ***(Douglas et al. 2005; Sergeeva et al. 2018)***. Using the Matlab program, 0.2 cycle/degree (15 sec per direction of rotation) was first used for mice to adapt this experiment, and 0.3, 0.35, 0.4, 0.43, 0.45, 0.47, 0.5 and 0.55 c/deg (30 sec per direction of rotation) were used in the following recordings. Mouse behaviors were analyzed in real time during the experiment and re-checked with video recordings. Finally, data for each mouse were determined by the minimal spatial frequency between left and right optomotor response.

### CTB Labelling of Optic Nerve

To label RGC axon terminals in mouse brain, RGC axons were anterogradely labeled by CTB (Cholera Toxin Subunit B) conjugated with Alexa Fluor™ 555 (Invitrogen, C34776) through intravitreal injection 48 hr before sacrifice. After PFA perfusion, the brains were fixed with 4% PFA in 0.1 M PB overnight, dehydrated with 15% sucrose and 30% sucrose in 0.1 M PB overnight at 4°C sequentially, embedded with O.C.T. for coronal section, and cryosectioned at 12 μm with Leica CM1950 Cryostat. After PBS washing, the sections were mounted with VECTASHIELD Antifade Mounting Medium with DAPI (Vector Laboratory). The images were captured on Tissue Genostics with identical settings for each group in the same experiment with the TissueFAXS 7.0 software.

### RNA immunoprecipitation and sequencing (RIP-Seq)

For RNA Immunoprecipitation (RIP) experiment, we used the EZ-Magna RIP^TM^ RNA-Binding Protein Immunoprecipitation Kit (Millipore) following the manual with minor modifications. Briefly, 1×10^7^ retina neurons were subjected to each 100 μl lysis buffer. The amount of YTHDF2 antibody (Proteintech, 24744-1-AP) and control IgG used for immunoprecipitation is 5 μg, respectively. Incubation was done overnight at 4 °C. After quality control monitoring using Agilent 2100, 100 ng RNA of input and elutes after RIP were used to generate the library using the TruSeq Stranded RNA Sample Preparation Kit (Illumina) and sequenced on the Illumina HiSeq 3000 platform (Jingneng, Shanghai, China). The filtered reads were mapped to the mouse reference genome (GRCm38) using STAR v2.5 ***(Dobin et al. 2013)*** with default parameters. The resulting bam files were fed to HTSeq tool ***(Anders et al. 2015)*** to count the number of RNA-seq reads, which was further normalized to calculate FPKM. To determine which gene is enriched, we computed the FPKM from RIP elute to input and any fold change greater than 2 (p value less than 0.05) was considered enriched. All enriched genes were used to do the Gene Ontology (GO) analyses. GO enrichment analysis was implemented by the GOseq R package, in which gene length bias was corrected. GO terms with corrected p value less than 0.05 were considered significantly enriched.

### MS analysis

E15.5 retinal neurons were cultured and infected with lenti viral *shYthdf2* or *shCtrl*. After puromycin (Sigma) selection, cells were washed with ice-cold PBS and then lysed with freshly prepared lysis buffer composed of 8 M urea (Sigma), 0.1 M HEPES (pH 7.4, Invitrogen), and protease inhibitors (Roche). The cell lysates were then ultrasonicated on ice and centrifuged at 10,000 × g for 10 min at 4 °C to discard the cell debris. Protein concentration was measured using the BCA Protein Assay Kit (Thermo Scientific). 100 μg of total protein for each group were reduced with 5 mM dithiothreitol (Sigma) for 30 min at 56 °C and then alkylated with 11 mM iodoacetamide (Sigma) for 15 min at RT in dark. After using 100 mM TEAB (Sigma) to dilute the urea concentration to less than 2 M in each sample, trypsin (Promega) was then added to digest the proteins overnight at 37 °C. Peptides were further desalted by Strata X C18 SPE column (Phenomenex) and labelled with TMT10plex Mass Tag Labelling kit (Thermo Scientific) according to the manufacturer’s instructions. Finally, the labeled peptides were subjected to HPLC fractionation and LC-MS/MS analysis. Proteins with fold changes greater than 1.3 and p values less than 0.05 were considered to be regulated by YTHDF2 KD with statistical significance.

### Anti-m^6^A Immunoprecipitation

Total retinal RNA was extracted from P0 WT mouse pups. Immunoprecipitation of m^6^A-modified transcripts was carried out with Magna MeRIP™ m^6^A Kit (Merck-Millipore, 17-10499) following the manual. m^6^A antibody (Synaptic Systems, 202003) and corresponding control IgG were used in this experiment. The RNA samples pulled down from the experiment were used for RT-qPCR.

### Acute ocular hypertension (AOH) model

Mice were anesthetized with 5% chloral hydrate in normal saline (10 μl/g) based on body weight and the Compound Tropicamide Eye Drops were used to scatter pupil. The anterior chamber was penetrated using the 32G × 1/2’’ needles (TSK) and filled with the BBS Sterile Irrigating Solution (Alcon) which was hung at a high position to provide proper pressure. Intraocular pressure was measured with the Tonolab tonometer (icare) for every 10 min and maintained at 85-90 mmHg for 1 hr. Levofloxacin hydrochloride was used after the operation and mice were revived in a 37°C environment. Retinas were analyzed for gene expression of YTHDF2 1 day after AOH, gene expression of *Hspa12a* and *Islr2* 3 days after AOH, dendritic complexity and RGC number 3-7 days after AOH.

### Statistical analysis

All experiments were conducted at a minimum of three independent biological replicates (two biological replicates for the RIP assay) or three mice/pups for each genotype/condition in the lab. Data are mean ± SEM. Statistical analysis was preformed using GraphPad Prism 7.0. When comparing the means of two groups, an unpaired or paired *t* test was performed on the basis of experimental design. The settings for all box and whisker plots are: 25th-75th percentiles (boxes), minimum and maximum (whiskers), and medians (horizontal lines). A *p* value less than 0.05 was considered as statistically significant: \**p* < 0.05, \*\**p* < 0.01, \*\*\**p* < 0.001, \*\*\*\**p* < 0.0001.

## Acknowledgements

We thank Ke Wang and Kwok-Fai So (Jinan University) for help on the optomotor response assay. We thank Mengqing Xiang and Suo Qiu (Zhongshan Ophthalmic Center, Sun Yat-sen University) for help on AAV experiments. We thank other members of Ji laboratory for technical support, helpful discussions and comments on the manuscript. This work was supported by National Natural Science Foundation of China (31871038 and 32170955 to S.-J.J.; 31922027 and 32170958 to B.P.), Shenzhen-Hong Kong Institute of Brain Science-Shenzhen Fundamental Research Institutions (2021SHIBS0002, 2019SHIBS0002), High-Level University Construction Fund for Department of Biology (internal grant no. G02226301), Science and Technology Innovation Commission of Shenzhen Municipal Government (ZDSYS20200811144002008), Program of Shanghai Subject Chief Scientist (21XD1420400), and the Innovative Research Team of High-Level Local University in Shanghai (B.P.).

## Author contributions

S.-J.J., F.N. and B.P. formulated the idea and designed the experiments; F.N. performed and analyzed most of the experiments; P.H., J.Z. and Y.S. carried out plasmid construction, AAV injection and imaging; L.Y. performed the RIP-seq experiment; J.Y. performed the MS experiment; M.Z., Y.S., B.Y. and C.L. provided technical help and helped with data analysis; F.N. and K.T. performed the AOH experiments under the supervision of B.P.; the optomotor code was written by B.P.; S.-J.J., F.N., L.Y., and J.Y. wrote the manuscript with inputs from other authors.

## Ethics

All experiments using mice were carried out following the animal protocols approved by the Laboratory Animal Welfare and Ethics Committee of Southern University of Science and Technology (approval numbers: SUSTC-JY2017004, SUSTC-JY2019081).

## Data availability statement

The RIP-seq data have been deposited to the Gene Expression Omnibus (GEO) with accession number GSE145390. The mass spectrometry proteomics data have been deposited to the ProteomeXchange Consortium via the PRIDE partner repository with the dataset identifier PXD017775.

**Supplementary file 1. List of YTHDF2 target mRNAs by anti YTHDF2 RIP-seq.**

**Supplementary file 2. Proteome of YTHDF2 knockdown vs control.**

**Supplementary file 3. Overlapping mRNA of Y2-RIP vs Y2-KD-MS.**

**Figure 1-source data 1. Source data for Figure 1B.**

(A) WB of anti YTHDF2 after KD of YTHDF2.

(B) WB of anti β-actin after KD of YTHDF2.

**Figure 1-source data 2. Source data for Figure 1B.**

Original file of the full raw unedited blot of anti YTHDF2 after KD of YTHDF2.

**Figure 1-source data 3. Source data for Figure 1B.**

Original file of the full raw unedited blot of anti β-actin after KD of YTHDF2.

**Figure 1-figure supplement 1-source data 1.**

Source data for Figure 1-figure supplement 1E,F.

(A) WB of anti YTHDF1 after KD of YTHDF1.

(B) WB of anti β-actin after KD of YTHDF1.

(C) WB of anti YTHDF3 after KD of YTHDF3.

(D) WB of anti β-actin after KD of YTHDF3.

**Figure 1-figure supplement 1-source data 2.**

Source data for Figure 1-figure supplement 1E.

Original file of the full raw unedited blot of anti YTHDF1 after KD of YTHDF1.

**Figure 1-figure supplement 1-source data 3.**

Source data for Figure 1-figure supplement 1E.

Original file of the full raw unedited blot of anti β-actin after KD of YTHDF1.

**Figure 1-figure supplement 1-source data 4.**

Source data for Figure 1-figure supplement 1F.

Original file of the full raw unedited blot of anti YTHDF3 after KD of YTHDF3.

**Figure 1-figure supplement 1-source data 5.**

Source data for Figure 1-figure supplement 1F.

Original file of the full raw unedited blot of anti β-actin after KD of YTHDF3.

---

**Figure 1–figure supplement 1.**
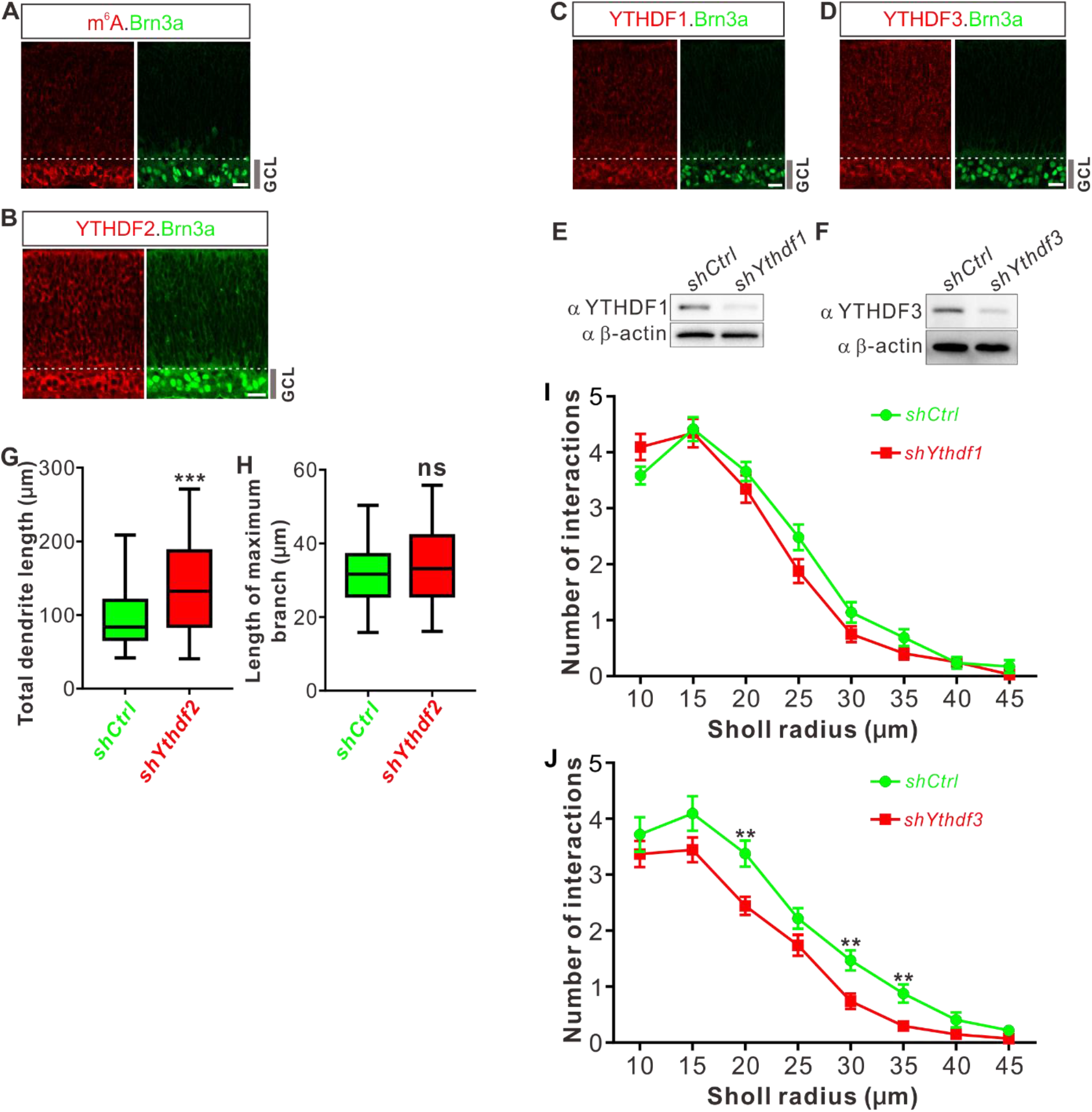
RGC have high level of m^6^A modification and strong expression of YTHDFs. (**A-D**) Representative confocal images showing high levels of m^6^A modification (**A**), and strong expressions of YTHDF2 (**B**), YTHDF1 (**C**), and YTHDF3 (**D**) in RGCs (marked by Brn3a) in P0 retina. Scale bars: 20 μm. (**E, F**) Western blotting (WB) confirming efficient knockdown (KD) of YTHDF1 and YTHDF3 in cultured RGCs using *shYthdf1* and *shYthdf3*, respectively. (**G, H**) Quantification of total length (**G**) and length of maximum branch (**H**) of RGC dendrites after YTHDF2 KD. Data are represented as box and whisker plots: *n* = 36 RGCs for *shCtrl*, *n* = 32 RGCs for *shYthdf2*; \*\*\**p* = 0.00062 for **G**; *p* = 0.22 for **H**; ns, not significant; by unpaired Student’s *t* test. (**I, J**) Quantification of dendrite branching using Sholl analysis after YTHDF1 KD (**I**) and YTHDF3 KD (**J**). Data are mean ± SEM. In **I**, *n* = 29 RGCs for *shCtrl*, *n* = 32 RGCs for *shYthdf1*, all not significant; in **J**, *n* = 32 RGCs for *shCtrl*, *n* = 27 RGCs for *shYthdf3*, \*\**p* = 0.0028 (20 μm), \*\**p* = 0.0028 (30 μm), \*\**p* = 0.0052 (35 μm); by unpaired Student’s *t* test.

**Figure 2–figure supplement 1.**
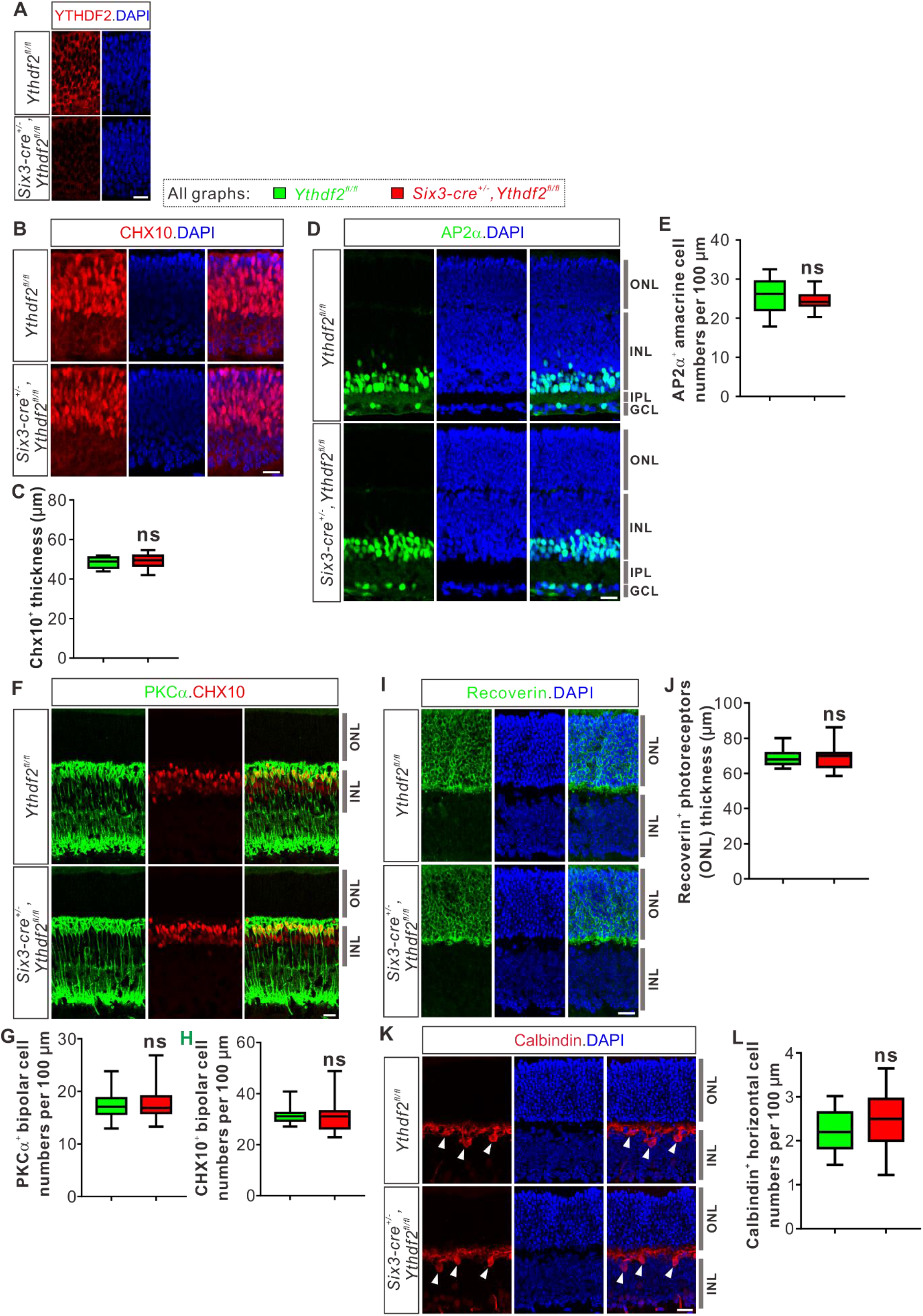
*Ythdf2* cKO does not change numbers of retinal progenitors, amacrine cells, bipolar cells, photoreceptors, or horizontal cells. (**A**) YTHDF2 protein was efficiently knocked out in the retinas of Six3-Cre-mediated *Ythdf2* cKO mice at E12.5. (B, C) Retinal progenitors not affected in *Ythdf2* cKO retina. CHX10 IF was used to label retinal progenitors at E15.5 (**B**). Thickness of CHX10^+^ retinal layer was quantified and showed no difference between the *Ythdf2* cKO retina (*n* = 10 sections) and their littermate controls (*n* = 19 sections) (**C**). (D, E) Amacrine cells not affected in the *Ythdf2* cKO retina. AP2α IF was used to mark amacrine cells in P6 retina (**D**). Numbers of AP2α^+^ amacrine cells per 100 μm of layer width in retina vertical sections were quantified and showed no difference between the *Ythdf2* cKO (*n* = 26 sections) and littermate controls (*n* = 26 sections) (E). ONL, outer nuclear layer; INL, inner nuclear layer; IPL, inner plexiform layer. (**F-H**) Bipolar cells not changed in the *Ythdf2* cKO retina. PKCα and CHX10 IF were used to label different bipolar cells in P15 retina (F). Numbers of PKCα^+^ and CHX10^+^ bipolar cells per 100 μm of layer width in retina vertical sections were quantified and showed no difference between the *Ythdf2* cKO (*n* = 18 sections for PKCα^+^ in G, *n* = 19 sections for CHX10^+^ in H) and littermate controls (*n* = 18 sections for PKCα^+^ in G, *n* = 19 sections for CHX10^+^ in H). (**I, J**) Photoreceptors not changed in the *Ythdf2* cKO retina. Recoverin IF was used to label photoreceptors in P20 retina (I). Thickness of Recoverin^+^ photoreceptor layer (e.g. ONL) in the retinal vertical sections was quantified and showed no difference between the *Ythdf2* cKO (*n* = 34 confocal fields) and littermate controls (*n* = 30 confocal fields) (J). (**K, L**) Horizontal cells not affected in the *Ythdf2* cKO retina. Calbindin IF was used to mark horizontal cells in P20 retina (arrowheads in K). Numbers of Calbindin^+^ horizontal cells per 100 μm of layer width in retina vertical sections were quantified and showed no difference between the *Ythdf2* cKO (*n* = 17 sections) and littermate controls (*n* = 17 sections) (L). All quantification data are represented as box and whisker plots: ns, not significant; *p* = 0.41 for C, *p* = 0.16 for E, *p* = 0.82 for G, *p* = 0.97 for H, *p* = 0.89 for J, *p* = 0.19 for L; by unpaired Student’s *t* test. Scale bars: 20 μm.

**Figure 3–figure supplement 1.**
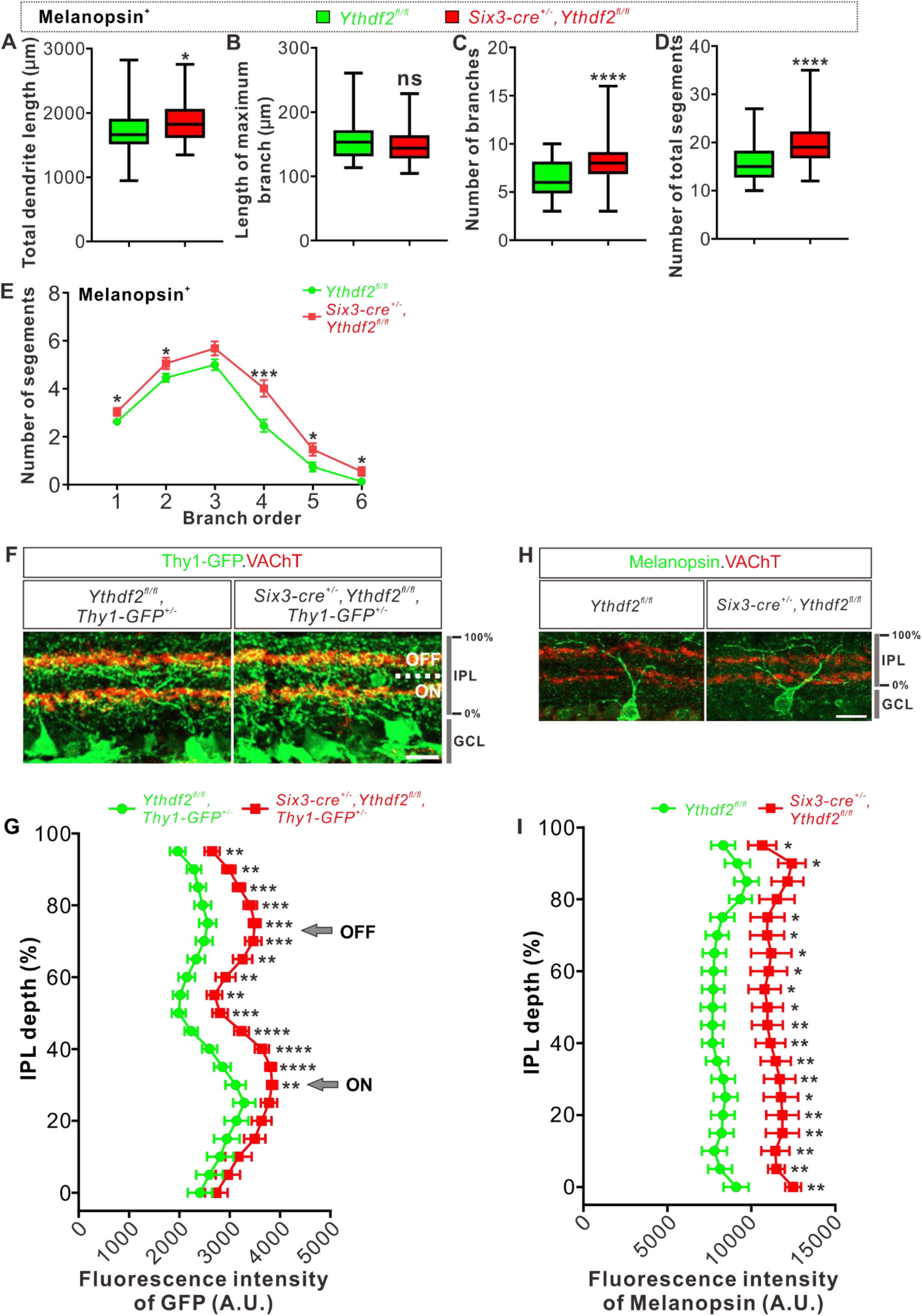
General dendrite density in IPL is increased without affecting sublaminar targeting. (**A-E**) Quantification of total length (**A**), length of maximum branch (**B**), branch numbers (**C**), number of total segments (**D**), and numbers of segments on each branch order (**E**) of melanopsin^+^ ipRGCs dendrites visualized by wholemount immunostaining of P20 retina using a melanopsin antibody in vivo (shown in Figure 3C). Data are represented as box and whisker plots in **A-D**: *n* = 58 RGCs for *Ythdf2^fl/fl^*, *n* = 51 RGCs for *Six3-cre^+/-^,Ythdf2^fl/fl^*; \**p* = 0.040 for **A**; *p* = 0.12 for **B**; \*\*\*\**p* = 1.39E-06 for **C**; \**p* = 7.89E-08 for **D**; ns, not significant. Data are mean ± SEM in **E:** \**p* = 0.038 (branch order 1), \**p* = 0.039 (branch order 2), \*\*\**p* = 0.00045 (branch order 4), \**p* = 0.026 (branch order 5), \**p* = 0.029 (branch order 6). All by unpaired Student’s *t* test. (**F**) Cross-sections of the IPL showing dendritic sublaminar patterning of Thy1-GFP^+^ RGCs in P20 control and *Ythdf2* cKO retina. ON and OFF refer to the ON-OFF bipartite divisions of the IPL marked by VAChT. Scale bar: 20 μm. (**G**) Quantification and distribution of GFP intensities from Thy1-GFP^+^ RGC dendrites through the depth of IPL shown in (**F**). GFP IF intensities are increased for the 30-95% depth of IPL in the *Ythdf2* cKO retina compared with their littermate controls, but the general patterning is similar between the two genotypes. Data are mean ± SEM (*n* = 13 sections for each genotype): \*\**p* = 0.0039 (95%), \*\**p* = 0.0014 (90%), \*\*\**p* = 0.00049 (85%), \*\*\**p* = 0.00020 (80%), \*\*\**p* = 0.00018 (75%), \*\*\**p* = 0.00036 (70%), \*\**p* = 0.0018 (65%), \*\**p* = 0.0067 (60%), \*\**p* = 0.0040 (55%), \*\*\**p* = 0.00057 (50%), \*\*\*\**p* = 3.48E-05 (45%), \*\*\*\**p* = 4.76E-05 (40%), \*\*\*\**p* = 6.85E-05 (35%), \*\**p* = 0.0034 (30%), by unpaired Student’s *t* test. Arrows indicate peaks of VAChT signals. (**H**) Cross-sections of the IPL showing dendritic sublaminar patterning of melanopsin^+^ ipRGCs in P20 control and *Ythdf2* cKO retina. Scale bar: 20 μm. (**I**) Quantification and distribution of melanopsin IF intensities from melanopsin^+^ ipRGC dendrites through the depth of IPL shown in (**H**). Melanopsin IF intensities are increased in the *Ythdf2* cKO retina compared with their littermate controls, but the general patterning is similar between the two genotypes. Data are mean ± SEM (*n* = 11 neurons for control, *n* = 8 neurons for *Ythdf2* cKO): \**p* = 0.049 (95%), \**p* = 0.010 (90%), \**p* = 0.039 (75%), \**p* = 0.022 (70%), \**p* = 0.019 (65%), \**p* = 0.018 (60%), \**p* = 0.016 (55%), \**p* = 0.013 (50%), \*\**p* = 0.0095 (45%), \*\**p* = 0.0044 (40%), \*\**p* = 0.0053 (35%), \*\**p* = 0.0091 (30%), \**p* = 0.014 (25%), \*\**p* = 0.0074 (20%), \*\**p* = 0.0071 (15%), \*\**p* = 0.0053 (10%), \*\**p* = 0.0025 (5%), \*\**p* = 0.0029 (0%), by unpaired Student’s *t* test.

**Figure 4–figure supplement 1.**
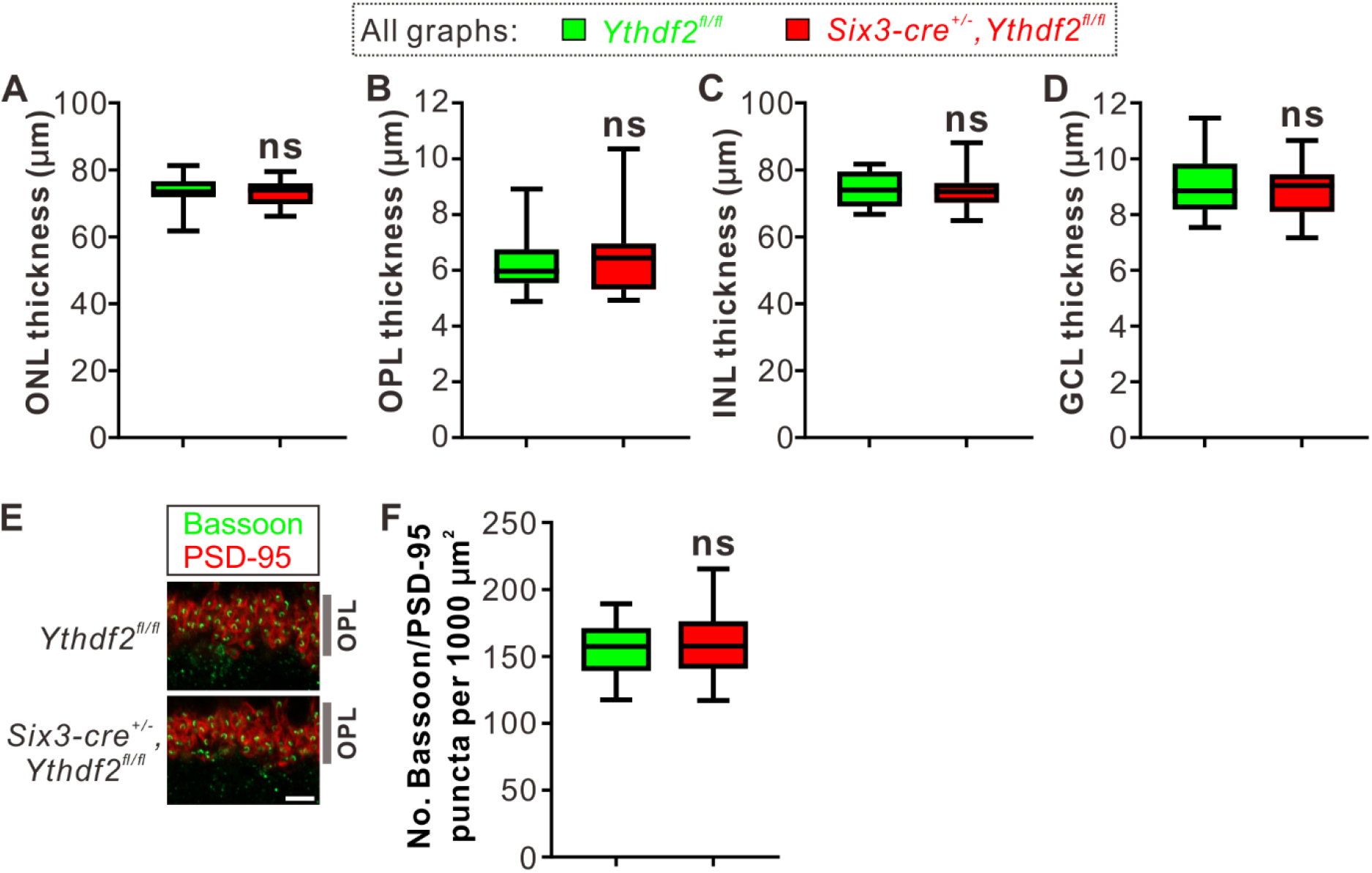
Thickness or synapse numbers in OPL shows no difference between the *Ythdf2* cKO and control retinas. (**A-D**) Quantification of the thickness of different layers by MAP2/DAPI IF in P6 *Ythdf2* cKO and control retinas as shown in Figure 4A. Data are represented as box and whisker plots: *n* = 14 sections for each genotype; *p* = 0.60 for ONL in **A**, *p* = 0.61 for OPL in **B**, *p* = 0.84 for INL in **C**, *p* = 0.62 for GCL in **D**; ns, not significant; by unpaired Student’s *t* test. (**E, F**) Representative confocal images showing the excitatory ribbon synapses labeled by co-localization of Bassoon (presynaptic) and PSD-95 (postsynaptic) in the OPL of P30 retina (**E**), which shows no difference between *Ythdf2* cKO and control. Quantification data are represented as box and whisker plots (**F**): *n* = 39 confocal fields for *Ythdf2^fl/fl^*, *n* = 36 confocal fields for *Six3-cre^+/-^,Ythdf2^fl/fl^*; *p* = 0.66; ns, not significant; by unpaired Student’s *t* test. Scale bar: 5 μm.

**Figure 5–figure supplement 1.**
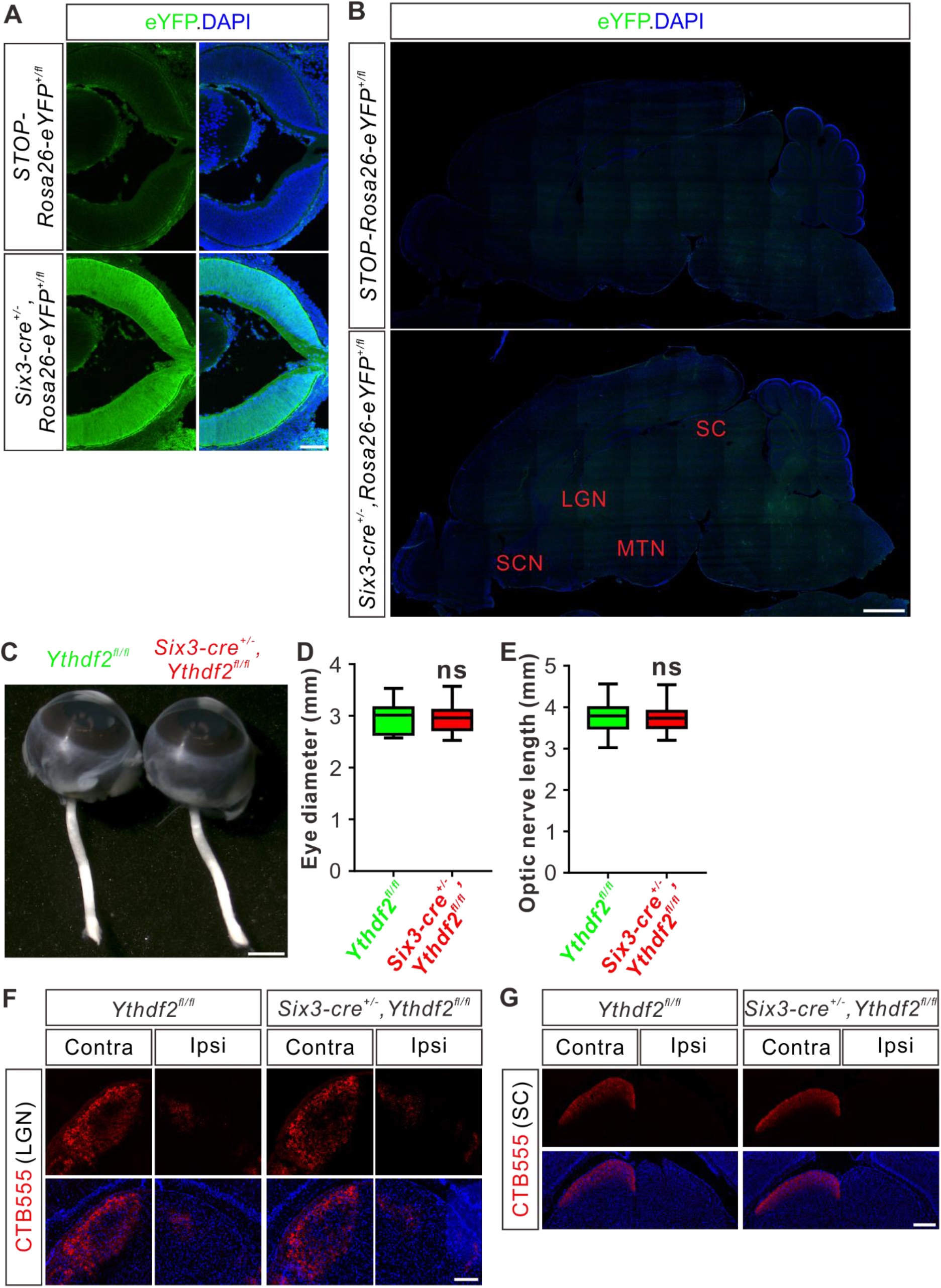
Guidance or central targeting of optic nerves is not affected in Six-Cre-mediated *Ythdf2* cKO. (**A**) Cross-sections of E14.5 retina showing strong expression of Six3-Cre using an eYFP reporter. Scale bar: 100 μm. (**B**) Sagittal sections of P10 brain showing negligible expression of Six3-Cre in the potential RGC target regions in the brain. SCN, suprachiasmatic nucleus; LGN, lateral geniculate nucleus; MTN, medial terminal nucleus; SC, superior colliculus. Scale bar: 1 mm. (**C-E**) Normal eye diameter and optic nerve length. Quantification data (**D, E**) are represented as box and whisker plots: *p* = 0.80 (*n* = 31 for each genotype in **D**); *p* = 0.99 (*n* = 31 for each genotype in **E**); ns, not significant; by unpaired Student’s *t* test. Scale bar: 1 mm. (**F, G**) Representative images of coronal sections through the LGN (**F**) and SC (**G**) after unilateral injection of CTB-Alexa Fluor 555 at P37 in *Ythdf2* cKO and control mice. Projections to the contralateral (Contra), ipsilateral (Ipsi) LGN and contralateral (Contra) SC are visible, which shows no difference between *Ythdf2* cKO and control mice. Scale bars: 100 μm (**F**) and 200 μm (**G**).

**Figure 7–figure supplement 1.**
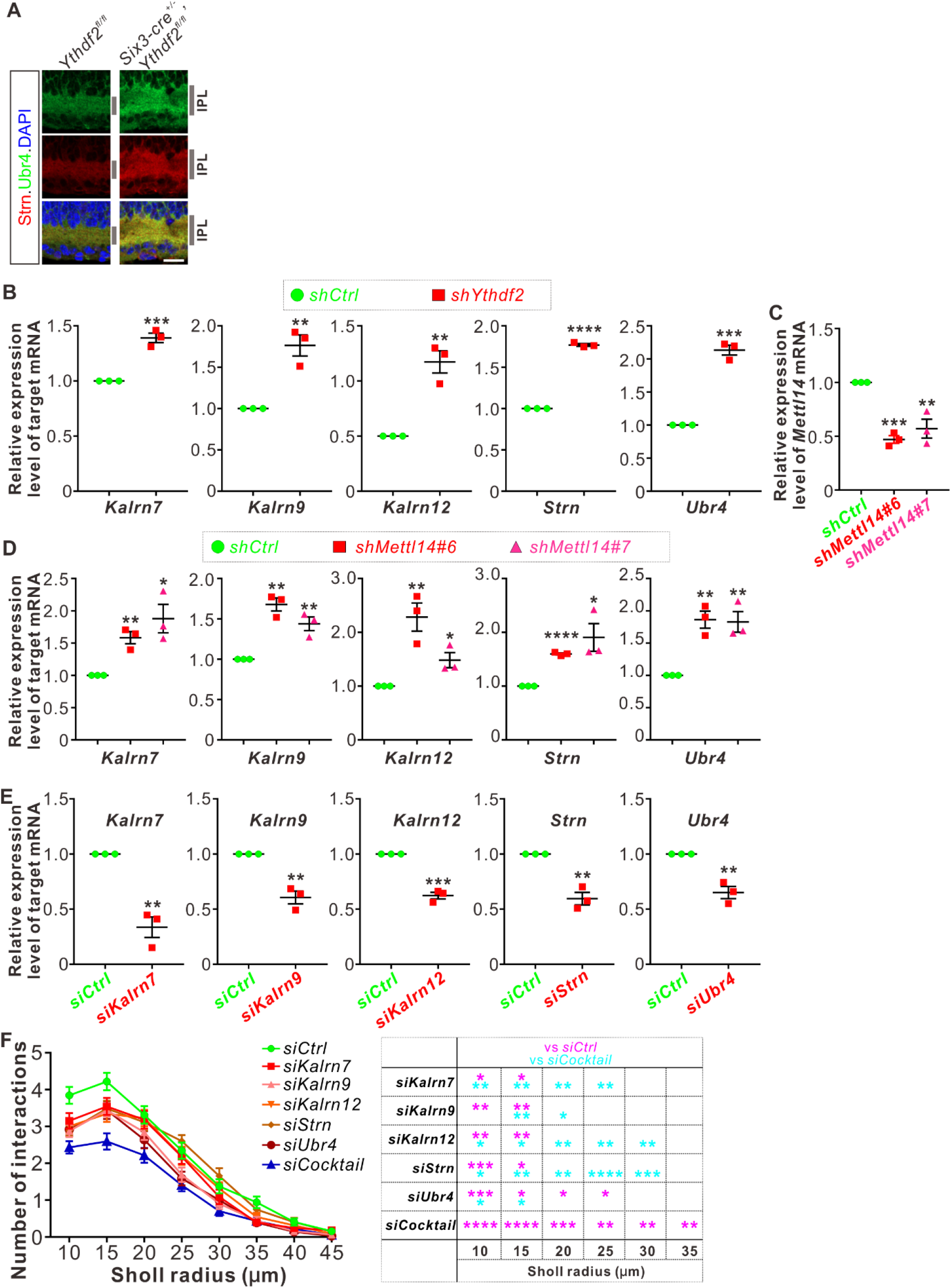
YTHDF2 target mRNAs were characterized and validated. (**A**) Upregulation of target mRNAs-encoding proteins Strn and Ubr4 in *Ythdf2* cKO retina in vivo. Enrichment and higher levels of these proteins were detected in the IPL of P6 *Ythdf2* cKO retina compared with control by IF. Scale bar: 20 μm. (**B**) Upregulation of target mRNA levels after YTHDF2 KD. RT-qPCR confirmed upregulation of the candidate target mRNAs after KD of YTHDF2 in cultured RGCs using *shYthdf2*. Data are mean ± SEM and are represented as dot plots (*n* = 3 replicates): \*\**\*p* = 0.00084 for *Kalrn7*; \**\*p* = 0.0039 for *Kalrn9*; \**\*p* = 0.0026 for *Kalrn12*; \*\*\**\*p* = 1.11E-06 for *Strn*; \*\**\*p* = 0.00011 for *Ubr4*; by unpaired Student’s *t* test. (**C**) Confirmation of METTL14 KD in cultured RGCs using *shMettl14* by RT-qPCR. Data are mean ± SEM and are represented as dot plots (*n* = 3 replicates): \*\*\**p* = 0.00012 (*shMettl14#6* vs *shCtrl*); \*\**p* = 0.0079 (*shMettl14#7* vs *shCtrl*); by unpaired Student’s *t* test. (**D**) Upregulation of target mRNA levels after METTL14 KD. RT-qPCR confirmed upregulation of the candidate target mRNAs after KD of METTL14 in cultured RGCs using *shMettl14*. Data are mean ± SEM and are represented as dot plots (*n* = 3 replicates): for *Kalrn7*, \*\**p* = 0.0034 (*shMettl14#6* vs *shCtrl*), \**p* = 0.016 (*shMettl14#7* vs *shCtrl*); for *Kalrn9*, \*\**p* = 0.0010 (*shMettl14#6* vs *shCtrl*), \*\**p* = 0.0067 (*shMettl14#7* vs *shCtrl*); for *Kalrn12*, \*\**p* = 0.0079 (*shMettl14#6* vs *shCtrl*), \**p* = 0.026 (*shMettl14#7* vs *shCtrl*); for *Strn*, \*\*\*\**p* = 5.45E-06 (*shMettl14#6* vs *shCtrl*), \**p* = 0.025 (*shMettl14#7* vs *shCtrl*); for *Ubr4*, \*\**p* = 0.0029 (*shMettl14#6* vs *shCtrl*), \*\**p* = 0.0066 (*shMettl14#7* vs *shCtrl*); by unpaired Student’s *t* test. (**E**) Confirmation of KD by siRNAs against target mRNAs. Data are mean ± SEM and are represented as dot plots (*n* = 3 replicates): \*\**p* = 0.0020 for *Kalrn7*; \*\**p* = 0.0025 for *Kalrn9*; \*\*\**p* = 0.00020 for *Kalrn12*; \*\**p* = 0.0021 for *Strn*; \*\**p* = 0.0033 for *Ubr4*; by unpaired Student’s *t* test. (**F**) KD of target mRNAs all together using a siRNA cocktail causing further decrease of dendrite branching of cultured RGCs compared with single siRNA against each target mRNA. Data are mean ± SEM: *n* = 32 RGCs for *siCtrl*, *n* = 33 RGCs for *siKalrn7*, *n* = 32 RGCs for *siKalrn9*, *n* = 35 RGCs for *siKalrn12*, *n* = 35 RGCs for *siStrn*, *n* = 36 RGCs for *siUbr4*, *n* = 36 RGCs for *siCocktail*. *siKalrn7* vs *siCtrl*: *\*p* = 0.031 (10 μm), \**p* = 0.046 (15 μm); *siKalrn9* vs *siCtrl*: \**\*p* = 0.0011 (10 μm), \*\**p* = 0.0090 (15 μm); *siKalrn12* vs *siCtrl*: \**\*p* = 0.0061 (10 μm), \*\**p* = 0.0086 (15 μm); *siStrn* vs *siCtrl*: \*\**\*p* = 0.00056 (10 μm), \**p* = 0.025 (15 μm); *siUbr4* vs *siCtrl*: \*\**p* = 0.0018 (10 μm), \**p* = 0.026 (15 μm), *\*p* = 0.048 (20 μm), *\*p* = 0.011 (25 μm); *siCocktail* vs *siCtrl*: \*\*\*\**p* = 3.44E-06 (10 μm), \*\*\*\**p* = 4.07E-06 (15 μm), \*\**\*p* = 0.00077 (20 μm), \**\*p* = 0.0010 (25 μm), \**\*p* = 0.0049 (30 μm), \**\*p* = 0.0094 (35 μm). *siKalrn7* vs *siCocktail*: \**\*p* = 0.0092 (10 μm), \*\**p* = 0.0040 (15 μm), \**\*p* = 0.0028 (20 μm), \**\*p* = 0.0034 (25 μm); *siKalrn9* vs *siCocktail*: \**\*p* = 0.0042 (15 μm), \**p* = 0.034 (20 μm); *siKalrn12* vs *siCocktail*: \**p* = 0.029 (10 μm), \**p* = 0.019 (15 μm), \*\**p* = 0.0014 (20 μm), \**\*p* = 0.0091 (25 μm), \**\*p* = 0.0063 (30 μm); *siStrn* vs *siCocktail*: \**p* = 0.043 (10 μm), \*\**p* = 0.0051 (15 μm), \*\**p* = 0.0045 (20 μm), \*\*\**\*p* = 3.79E-06 (25 μm), \*\**\*p* = 0.00022 (30 μm); *siUbr4* vs *siCocktail*: \**p* = 0.049 (10 μm), \**p* = 0.011 (15 μm). All by unpaired Student’s *t* test.

**Figure 8–figure supplement 1.**
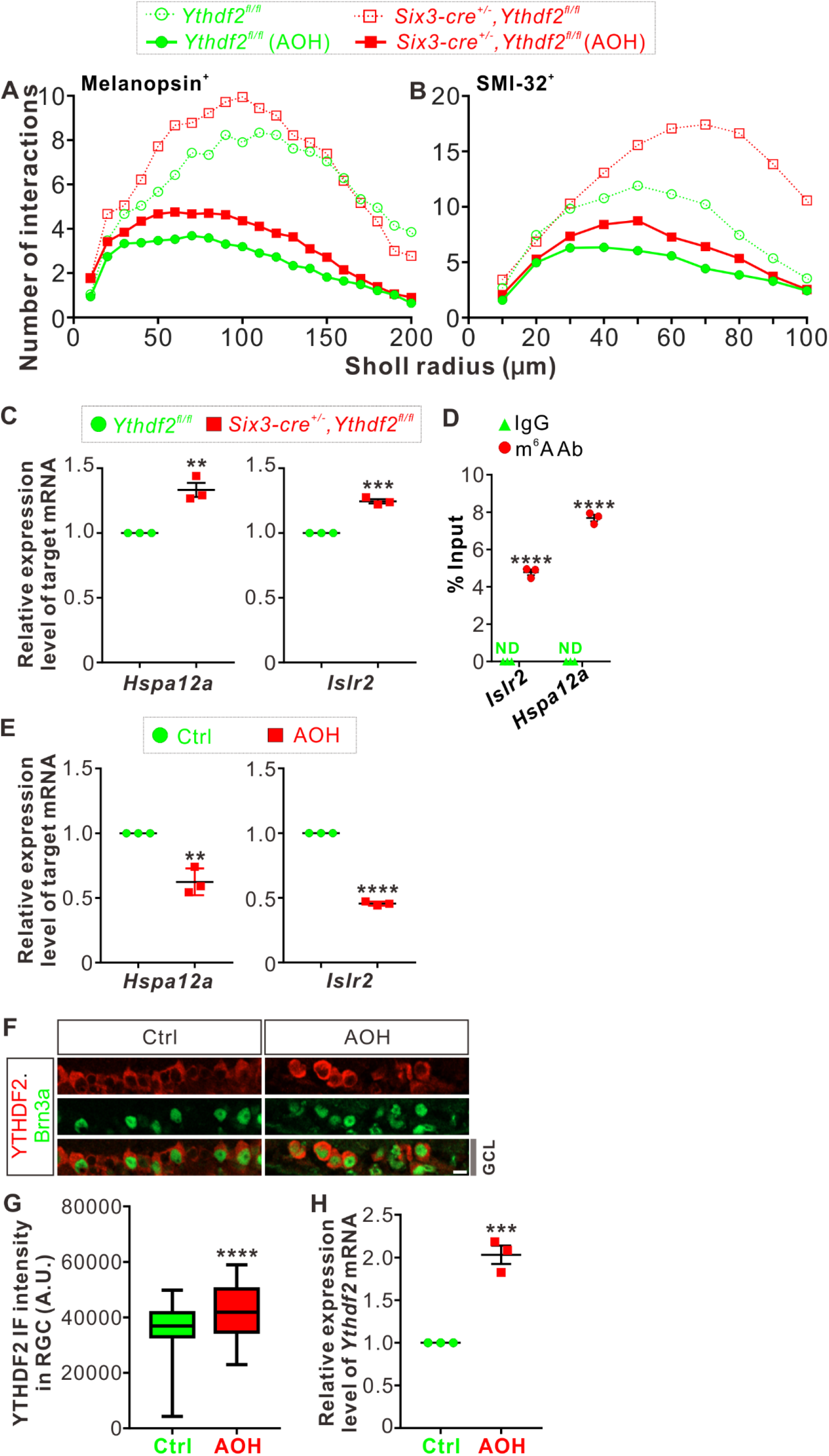
*Hspa12a* and *Islr2* are two target mRNAs of YTHDF2 in adult retina. (**A, B**) The curves from Figure 3D,F and Figure 8A,B were plotted together for easy comparison. The error bars and the asterisks were removed from these graphs for easy reading and these information can still be seen in Figure 3D,F and Figure 8A,B. (**C**) Upregulation of YTHDF2 target mRNA *Hspa12a* and *Islr2* in adult *Ythdf2* cKO retina compared with control by RT-qPCR. Data are mean ± SEM and are represented as dot plots (*n* = 3 replicates): \*\**p* = 0.0035 for *Hspa12a*; \*\*\**p* = 0.00012 for *Islr2*; by unpaired Student’s *t* test. (**D**) Verification of m^6^A modification of *Hspa12a* and *Islr2* mRNAs by anti-m^6^A pulldown followed by RT-qPCR. ND, not detected. Data are mean ± SEM and are represented as dot plots (*n* = 3 replicates): \*\*\*\**p* = 7.07E-06 for *Islr2*; \*\*\*\**p* = 1.55E-06 for *Hspa12a*; by unpaired Student’s *t* test. (**E**) Downregulation of *Hspa12a* and *Islr2* mRNA levels in retina 3 days after AOH. Data are mean ± SEM and are represented as dot plots (*n* = 3 replicates): \*\**p* = 0.0032 for *Hspa12a*; \*\*\*\**p* = 5.41E-07 for *Islr2*; by unpaired Student’s *t* test. (**F, G**) Cross-sections of retina showing increased YTHDF2 expression in Brn3a^+^ RGCs by IF. AOH was performed using P60 mice, and retinas were collected 1 day after AOH for analysis. Quantification data of YTHDF2 IF were represented as box and whisker plots (**G**): \*\*\**\*p* = 1.13E-06 (*n =* 110 RGCs for each condition); by unpaired Student’s *t* test. Scale bar: 10 μm. (**H**) Upregulation of *Ythdf2* mRNA level after AOH. Data are mean ± SEM and are represented as dot plots (*n =* 3 replicates): \*\**\*p* = 0.00066; by unpaired Student’s *t* test.

